# A null model of the mouse whole-neocortex micro-connectome

**DOI:** 10.1101/548735

**Authors:** Michael W Reimann, Michael Gevaert, Ying Shi, Huanxiang Lu, Henry Markram, Eilif Muller

## Abstract

Connectomics, the study of the structure of networks of synaptically connected neurons, is one of the most important frontiers of neuroscience. Great advances are being made on the level of macro- and meso-scale connectomics, that is the study of how and which populations of neurons are wired together by tracing axons of anatomically and genetically defined neurons throughout the brain. Similarly, the use of electron-microscopy and statistical connectome models has improved our understanding of micro-connectomics, that is the study of connectivity patterns between individual neurons. We have combined these two complementary views of connectomics to build a first draft statistical model of the neuron-to-neuron micro-connectome of a whole mouse neocortex. We combined available data on region-to-region connectivity and individual whole-brain axon reconstructions to model in addition to the meso-scale trends also the innervation of individual neurons by individual axons, within and across regions. This process revealed a novel targeting principle that allowed us to predict the innervation logic of individual axons from meso-scale data. The resulting micro-connectome of 10 million neurons and 88 billion synapses recreates biological trends of targeting on the macro-meso- and micro-scale, i.e. targeting of brain regions, domains and layers within a brain region down to individual neurons. This openly accessible connectome can serve as a powerful null model to compare experimental findings to and as a substrate for whole-brain simulations of detailed neural networks.

## 2 Introduction

The study of connectomics has to date largely taken place on two separate levels with disjunct methods and results: macro-connectomics, studying the structure and strength of long-range projections between brain regions, and micro-connectomics, studying the topology of individual neuron-to-neuron connectivity within a region.

In macro-connectomics, the absence or presence and strength of projections between brain regions are measured using for example histological pathway tracing, retrograde (Markov et al., 2014; Gămănuţ et al., 2018) or anterograde (Harris et al., 2018) tracers or MR diffusion tractography (Calabrese et al., 2015; van den Heuvel et al., 2015). While recent advances made it possible to turn such data into connectome models with a resolution of 100*µm* (Knox et al., 2018), this is still far away from single neuron resolution.

In micro-connectomics, two complementary approaches prevail: continuously refined stochastic models and direct measures of synaptic connectivity using for example electron-microscopy. The first uses biological findings to formulate principles that rule out certain classes of wiring diagrams and prescribe probabilities to the remaining ones, then instantiating a number of diagrams according to that stochastic model for study or simulation.

Such principles comprise among others distance-dependent connectivity (Petersen and Sakmann, 2000; Feldmeyer et al., 2002; Feldmeyer, 2006), connectivity depending on axons and dendrites reaching each other (“Peters’ Rule”, Peters et al. (1979); Hellwig (2000)), connectivity depending on neuronal types (Bannister, 2005; Jiang et al., 2015; Markram et al., 2015), limited axonal space for synapse formation (Reimann et al., 2015) and stable connections requiring multiple synapses (Reimann et al., 2015; Markram et al., 2015). The principles can drastically shrink the space of possible wiring diagrams (Reimann et al., 2017) and in the remaining diagrams, biologically identified connectivity trends emerge. Yet, until all principles of biological connectivity have been correctly identified, the space of possible wiring diagrams will contain a large number of unbiological ones. On the other hand, the availability of a continuous space of possible diagrams enables simulation studies of structural synaptic plasticity (Holtmaat and Svoboda, 2009).

With electron microscopy, snapshots of individual wiring diagrams are taken. The approach measures connectivity by reconstructing neurons and their synapses in a volume of interest (Denk and Horstmann, 2004; Briggman and Denk, 2006; Chklovskii et al., 2010; Kleinfeld et al., 2011; Helmstaedter et al., 2013; Kasthuri et al., 2015; Lee et al., 2016). While the resulting connectivity is guaranteed to be biological, the destructive nature of the approach means one cannot consider its evolution through plasticity. And published reconstructed volumes at this point only contain incomplete dendritic trees and therefore incomplete connectivity.

Both approaches to micro-connectomics are limited to local connectivity within a brain region, or along individual, important pathways, such as hippocampal CA3 to CA1, and consequently ignore a substantial amount of external synaptic input onto neurons. Yet, models based on only local connectivity have led to important insights (Markram et al., 2015). Macro-connectomics on the other hand lack single-neuron resolution, but have been successfully employed, for example in mean-field brain models (Becker et al., 2015; Roy et al., 2014; Sigala et al., 2014).

To gain a full understanding of, for example the role of an individual neuron or small groups of neurons in a given behavior, we will have to integrate the advantages of both scales: single neuron resolution on a whole-brain or at least whole-neocortex level. This has been recognized before (Bota et al., 2015), but steps towards this goal have until now remained limited. At this point, electron-microscopic reconstructions at that scale are not viable, leaving only statistical approaches to dense micro-connectivity, based on identifying biological principles in the data. Scaling it up to a whole-neocortex level will amplify the uncertainty about the biological accuracy of the results, as many of the resulting connections will be between rarely studied brain regions with little available biological data. Nevertheless, it can serve as a first draft micro-connectome defining a null model to compare and evaluate future findings against. It will also allow us to perform full-neocortex simulations at cellular resolution to gain insights, as to which brain function can or cannot be explained with a given connectome.

We have completed such a first-draft connectome of mouse neocortex by using an improved version of our previously published circuit building pipeline (Markram et al., 2015). It has been improved to place neurons in brain-atlas defined 3d spaces instead of hexagonal prisms, taking into account the geometry and cellular composition of individual brain regions. The composition was based on data from (Erö et al., 2018). The local connectivity, i.e. synaptic connections within a brain region, were placed based on axo-dendritic appositions and biological principles, as previously published (Reimann et al., 2015). However, this did not include long-range connections between brain regions, especially the ones formed via projections along the white matter. We therefore set out to identify possible principles, hypotheses of rules constraining the long-range connectivity, and develop stochastic methods to instantiate micro-connectomes fulfilling them.

A first constraint was given by data on macro- or meso-scale connectivity, which is often reported as a region-to-region connection matrix, yielding a measure proportional to the total number of synapses forming a projection between pairs of brain regions (Scannell et al., 1995; Markov et al., 2014; Zingg et al., 2014; Bota et al., 2015). We used for this purpose a recently published meso-scale mouse brain connectome (Harris et al., 2018). This dataset splits the mouse neocortex into 86 separate regions (43 per hemisphere) and further splits each region when considered as a source of a projection into five individual *projection classes*, by layer or pathway (Layer 23IT, Layer 4IT, Layer 5IT, Layer 5PT, Layer 6CT). IT refers to intratelencephalic projections, targeting ipsi- and contralateral cortex and striatum; PT refers to pyramidal tract projections, predominantly targeting subcortical structures, but also ipsilateral cortex; CT refers to cortico-thalamic projections. From here on we will leave out this additional distinction for projections from layers 2/3, 4 and 6, where only one class is specified in the data of Harris et al. (2018).

As the data in Harris et al. (2018) is focused on the right hemisphere, we assumed connectivity to be symmetrical between hemispheres to be able to model both of them. This lead to 5 (projection classes) ×43 (source regions) ×86 (ipsi- and contra-lateral target regions) potential projections parameterized in terms of their strength by the data. However, we considered the 5 × 43 ipsi-lateral projections within the same region to be local connectivity, which we instead derived with our established approach (Reimann et al., 2015). A number of regions also lack layer 4, rendering projections in that projection class void.

We further constrained the spatial structure of each projection within the target region. Along the vertical axis (orthogonal to layer boundaries) this was achieved by assigning a layer profile to each projection, as provided by Harris et al. (2018). Along the horizontal axes, we assumed a generalized topographical mapping between regions, implemented by defining local coordinate systems in the source and target regions and mapping points at corresponding coordinates to each other. These coordinate systems were parameterized using a voxelized (resolution 100*µm*) version of the data provided by Knox et al. (2018).

As a final constraint we applied rules on the number and identity of brain regions innervated by individual neurons in a given source region. To this end we analyzed the brain regions innervated by individual in-vivo reconstructions of whole-brain axons in a published dataset (Mouse-Light project at Janelia, mouselight.janelia.org, Gerfen et al. (2018)). Based on the analysis, we conceptualized and parameterized a decision tree of long-range axon targeting that reproduced the targeting rules found in the in-vivo data. This approach could be generalized to other brain regions for which few or no axonal reconstructions are available.

Finally, we implemented a stochastic algorithm that connected morphologically detailed neurons in a 3d-volume representing the entire mouse neocortex. Synapses were placed onto the dendrites of target neurons according to all the derived constraints by a modified version of a previously used algorithm (Markram et al., 2015).

## 3 Results

We built a morphologically detailed model of mouse neocortex using a scaled-up version of our previously published circuit building pipeline (Markram et al., 2015). Briefly, the workflow placed around 10 million neuron reconstructions in a 3d space representing the entirety of a mouse neocortex. Neuron densities and excitatory to inhibitory ratios at each location were taken from a voxelized brain atlas (Erö et al., 2018), that is consistent with version 3 of the brain parcellation of the Allen Brain Atlas (Dong, 2008; noa, 2017). The composition in terms of morphological neuron types was as in (Markram et al., 2015). Reconstructed morphologies were placed in the volume according to densities for individual, morphologically defined subtypes and correctly oriented with respect to layer boundaries.

For simplicity, we made a strict distinction between local and long-range connectivity, defining local connectivity to comprise any connection where source and target neuron were in the same brain region according to the parcellation in Harris et al. (2018), and long-range connectivity to make up all other connections. We therefore split the model according to that parcellation scheme and derived local connectivity for neurons within each subset as previously published (Reimann et al., 2015).

### 3.1 Constraining the anatomical strengths of projections

For long-range connectivity, we handled each combination of a projection class (Layer 23, Layer 4, Layer 5IT, Layer 5PT, Layer 6), a source region, and a target region as a conceptually separate projection. In this view, non-existing projections are considered projections with zero synapses. As a first constraint, we determined the average volumetric density of synapses in each projection using published data (Harris et al., 2018), using a programmatic interface provided by the authors. Two further steps were required to apply their data: scaling from projection strength to synapse density, and splitting into densities for individual projection classes.

Concerning the first step, their published “connection density” data is described as the total amount of signal from labelled axons in a region divided by the volume of the region, i.e. it can be considered proportional to the mean volumetric density of projection axons. Assuming a uniform mean density of synapses on axons across projections, the volumetric synapse density is simply a scaled version of the “connection density”. We calculated a scaling factor such that the resulting total synapse density in ipsi- and contra-lateral projections matches previously published results (Schüz and Palm, 1989). From their measured average synapse density (0.72*µm^−^*^3^) we subtracted the synapses we predicted in local connectivity within a region. While a part of the remaining synapses is formed by projections from hippocampus and extra-cortical structures, their total number is unclear, but likely comparatively small. For example, the density of synapses in the prominent pathway from VPM into the barrel field (Meyer et al., 2010), when averaged over the whole cortical depth, is only about 1.5% of the average total density. For now we left no explicit space for synapses from such projections due to the difficulties in parameterizing it for all potential sources.

The resulting matrix of synapse densities was then further broken up into matrices for different projection classes by first averaging the reported strengths of projection classes over *modules* of several contiguous brain regions (see Table S1), and then using that information to generate scaled versions of the matrix (see methods). The result is a prediction of the mean volumetric synapse densities from the bottom of layer 6 to the top of layer 1 for all projections (Fig. 1). To reduce the computational demand of the following steps, we determined a minimal projection strength and removed projections weaker than the cutoff. The cutoff was calculated as 0.0006*µm^−^*^3^, such that less than 5% of projection synapses would be lost.

**Figure 1:**
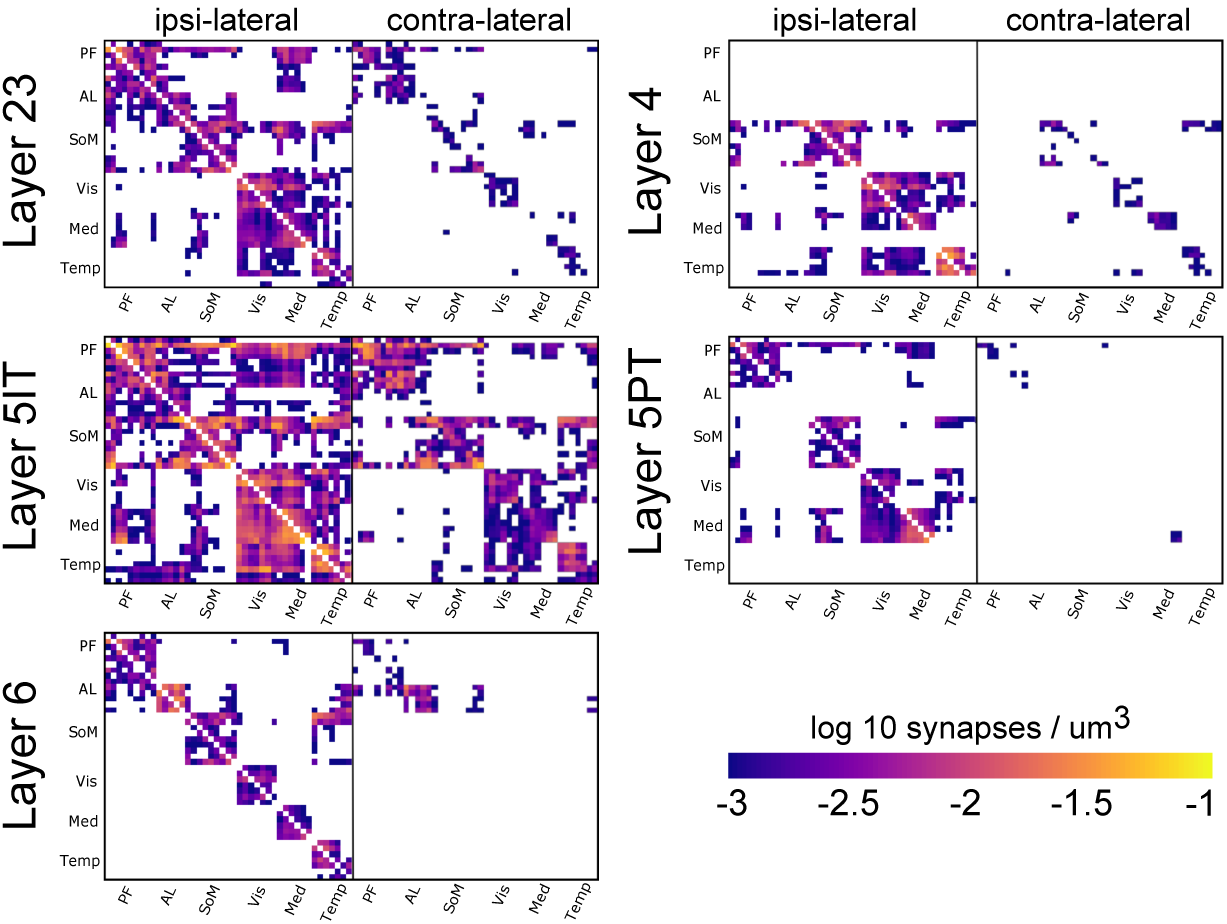
Predicted mean synapse density in target regions for all projections. Modules are labeled: PF: prefrontal, AL: anterolateral, SoM: somatomotor, Vis: visual, Med: medial, Temp: temporal. Exact order of brain regions and assignment to modules as in Harris *et al.*, 2018, also listed in Table 1. White regions indicate no projections placed for that combination of source and target region.

### 3.2 Constraining layer profiles

So far, we have constrained density and consequently the total number of synapses formed by each individual projection. This reproduces the spatial structure of projections on the macro-scale. However, it is likely that there is also spatial structure within a projection, on the meso- or micro-scale. One such micro-structure, acting along the vertical axis is a distinct targeting of specific layers (Van Essen et al., 1992). To constrain the layer profiles of projections, we once more tended to the data published in Harris et al. (2018). The authors provide extensive data on layer profiles, measured hundreds of them and then clustered them into six prototype profiles. We decided to follow this classification and assign one of the prototype profiles to each projection.

Harris et al. (2018) already measured the relative frequencies of their prototypical layer profiles for individual projection classes (their Figure 5o) and for individual source modules, within and across modules (their Figure 8c,d). They also classified profiles as belonging to feed-forward or feed-back projections. We combined the constraints by first calculating which layer profiles are over- or under-expressed between pairs of modules, relative to the base profile frequencies for projection classes (see Methods). We then classified each projection as feed-forward or feed-back, based on the hierarchical position of the participating regions, and cut the assumed frequencies of profiles belonging to the other type in half. Finally, we picked for each combination of projection class, source and target region the layer profile with the highest derived frequency.

We chose to pick the single most likely profile for each projection and ignore the others, as mixing several profiles would have diminished their sharp, distinguishable peaks and troughs. The approach resulted in a prediction, where each profile is used for between 10 and 20% of the projections (Fig. 2A). Based on the prediction we calculated the resulting relative frequencies of layer profiles per module and per projection class and compared them against the data (Fig. 2B). We found that in spite of the simplifying step of picking only the most likely profile, the trends in the data were well preserved, although the peaks and troughs were more exaggerated in the model.

**Figure 2:**
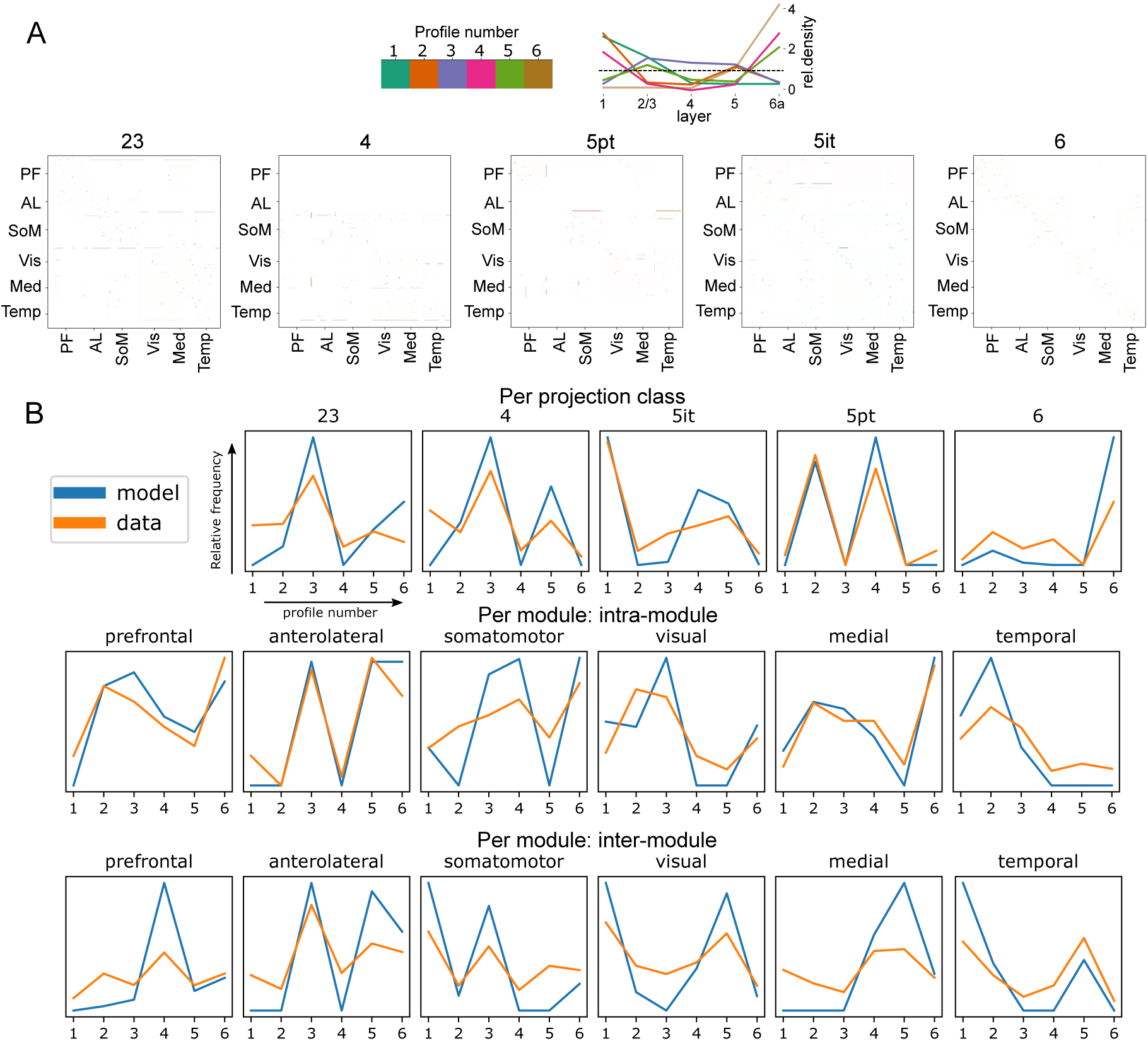
A: Predicted layer profiles for all projection classes. Exact order of brain regions and assignment to modules as in Harris et al., 2018, also listed in Table S1. B: Comparing resulting layer profile frequencies for projection classes and modules to the data of Harris et al., 2018

#### 3.2.1 Validating predicted layer profiles against raw data

We have demonstrated that our simplified predictions recreate the tendencies demonstrated in Harris et al. (2018), but the question remains, how do they compare against the raw biological data? As we moved through two consecutive simplifications - from the raw data to six prototypical profiles and from six profiles to a single profile per projection - how much biological detail was lost?

To address this question, we collected all available layer profiles for all projections from a voxelized version of the mouse meso-connectome model (Knox et al., 2018) and compared them to our single prediction. This data yields the connection strength measured with individual tracer experiments in voxels with a resolution of 100*µm*^3^. For each experiment, we calculated a profile of the connection strengths in each layer of a region, relative to the mean across all layers. As a representative example, Figure 3 depicts the model for projections from MOs (blue line) and the data from individual experiments (grey lines). We see an overall fair match between simplified prediction and data, albeit with some errors. For example, the data for 5PT projections show very shallow profiles in 4 regions that were not predicted. For 5IT projections to visual regions the data flattens out in layer 1 instead of peaking, although this may be partly artificial, because the data resolution of 100*µm*^3^ is close to the width of layer 1, leading to unreliable sampling.

**Figure 3:**
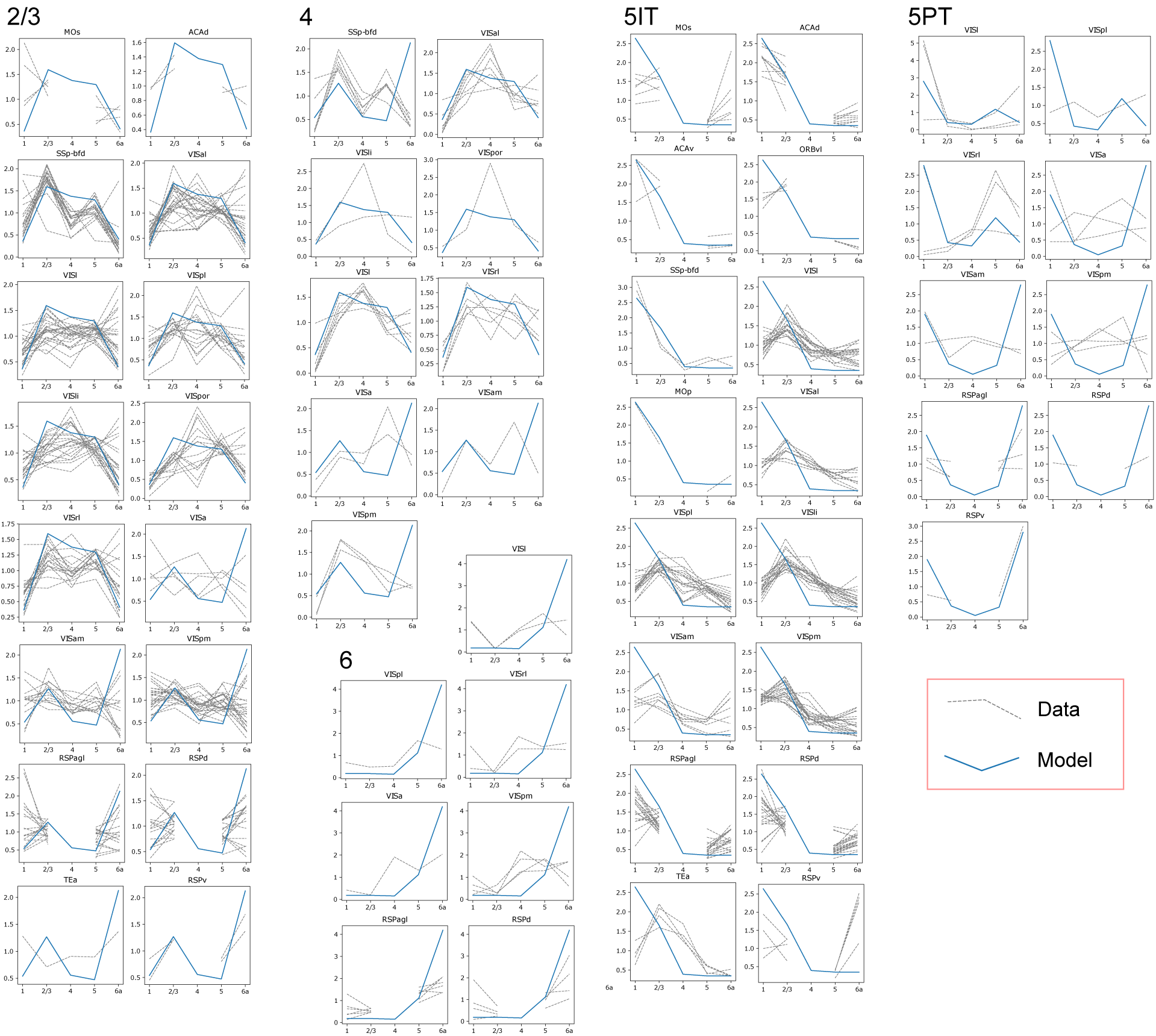
Comparing raw layer profiles from individual experiments in the voxelized projection model (grey, Knox et al., 2018) to our model prediction in all regions due to projections from neurons in MOs. Only data where the projection strength in a region is more than 1% of the total strength in all regions is shown.

Overall, we find a substantial degree of variability in the biological data, especially for projections from layer 2/3. For example, the density in layer 4 of VISpor due to projections from layer 2/3 of MOs varies between 0.2 and 2.5 times mean. As such, we evaluated the overall match of our predictions relative to the biological variability by calculating the deviation from the biological mean in multiples of the biological standard deviation (z-score, Figure 4 A-E). As a certain number of samples is required to estimate the biological variability, we limited this validation to projections where data from at least five experiments was available. Under the assumption of a gaussian profile, data randomly sampled from the biological distribution would follow a standard normal distribution of z-scores (Figure 4, black dashed lines). We found that the bulk of our predictions fall within that distribution, although a significant number have a z-score exceeding 4 standard deviations, especially for projections from layer 4 (Figure 4 A-E). Yet, 75% of z-scores fall within two standard deviations (Figure 4 F).

**Figure 4:**
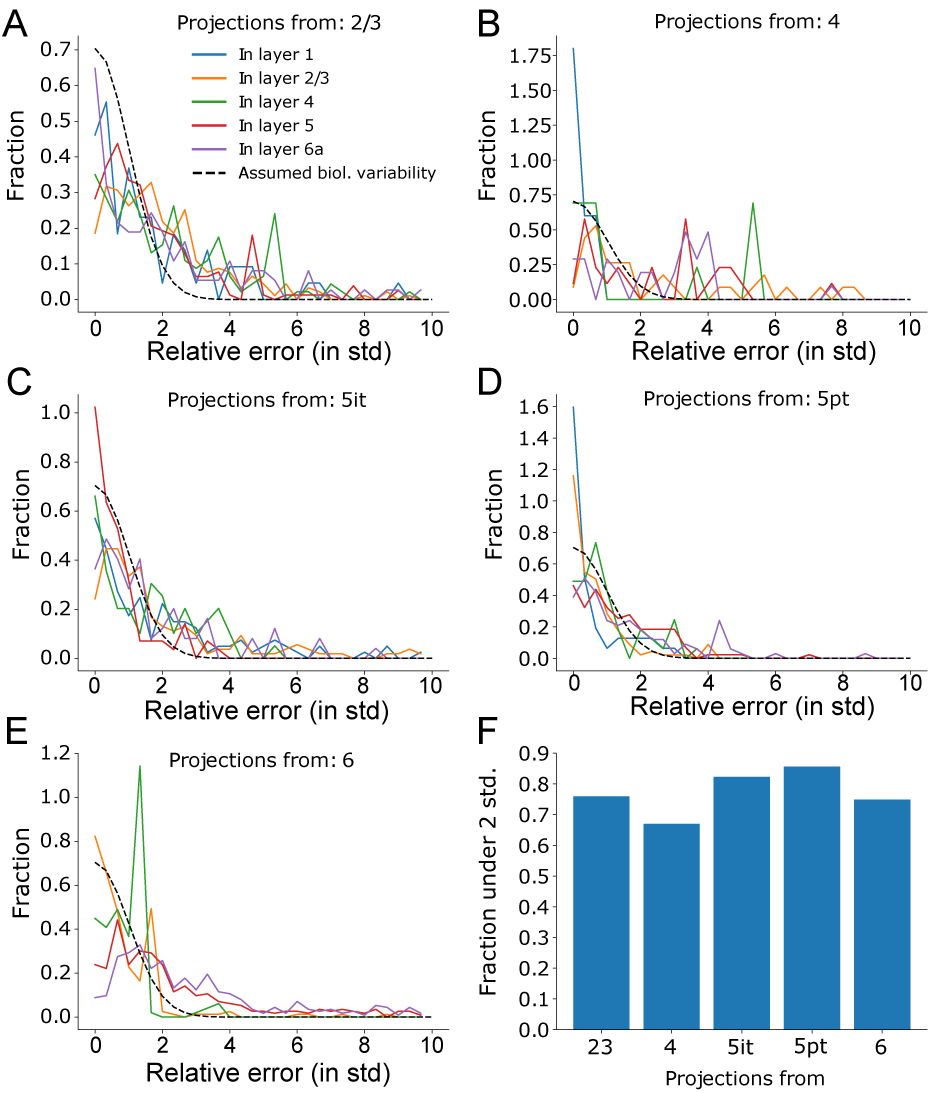
Validating the predicted layer profiles against raw data and the biological variability in it. A-E. Relative error of the predicted synapse densities in all layers. That is, the difference between prediction and the mean of the raw biological data, divided by the standard deviation of the biological data. A: For projections from L2/3, B: from L4, C: from L5IT, D: from L5PT, E: from L6. Dashed black lines indicate the biological variability of density under the assumption that it is gaussian distributed. We used only projections where more than 5 raw data points to establish the biological variability were available. F: Fraction of projections where with a relative error under two standard deviations for each source layer.

We conclude that the predicted layer profiles fall within the range of biological variability for most projections but do result in imperfect densities in individual cases. We judge this to be sufficient for a first draft null model of a white matter micro-connectome, but refinement should be attempted in the future, as more data, such as whole-brain axonal reconstructions become available.

### 3.3 Constraining the mapping of projections

The previous section constrained projection by imposing a spatial structure along the vertical axis, a layer profile. Yet, it is likely that there is also a structure along the other two spatial dimensions. That is, that neurons around a given point in the source region project not equally to all points in the target region, but with certain spatial preferences, which we assumed can be expressed by a *topographical mapping*. We defined a general topographical mapping between pairs of regions in a three step procedure: First, we projected 3d representations of the regions into 2d, preserving distances along the cortical surface (as in Harris et al. (2018)). This effectively collapsed the vertical axis, as we had constrained structure along that axis in the previous step. Next, we defined a local barycentric coordinate system in the 2d representation of the source region by selecting three points such that the resulting triangle covers most of the region. Finally, we defined a local barycentric coordinate system in the target region based on biological data on the modeled projection.

#### 3.3.1 Parameterizing the mapping of projections

To select appropriate points in the target region, we once more used the voxelized version of the mouse meso-connectome model (Knox et al., 2018). As each brain region comprised many voxels in the model, we could use this data to determine whether any given part of a brain region projected more strongly to some part of the target region than to other parts. This would indicate a structured, non-random mapping that we would have to recreate to preserve the biologically accurate cortical architecture.

As mentioned above, we first defined a *barycentric source coordinate system* in the 2d-projected source region. The points defining the coordinate system were first picked by maximizing the pairwise distances between them while staying inside the source region and then moved 25% towards the center of gravity of the region. We visualized the result by setting each of the red, green and blue color channels of an image of the source region to one of the three barycentric coordinates (Fig. 5A, *C_src_*). By extension, we also associated each voxel of the macro-connectome model (*x, y, z*) with a color (*B_x,y,z_*) by first projecting its center into the 2d plane, then looking up the barycentric coordinate. Next, we considered the strengths of projections from each source voxel and visualized the results by coloring each target pixel according to the product of the 2d-projected projection strength and the color associated with the source voxel:

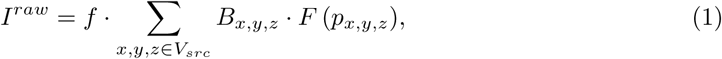

where *p_x,y,z_* refers to the voxelized projection strength from the voxel at *x, y, z*, *F* (*p_x,y,z_*) to its 2d projection and *f* to a scaling factor effectively deciding the overall lightness of the resulting image. The result is a two-dimensional image with three color channels 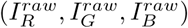. To more clearly reveal the structure of projections, we ignored source voxels associated with a color saturation below 0.5.

**Figure 5:**
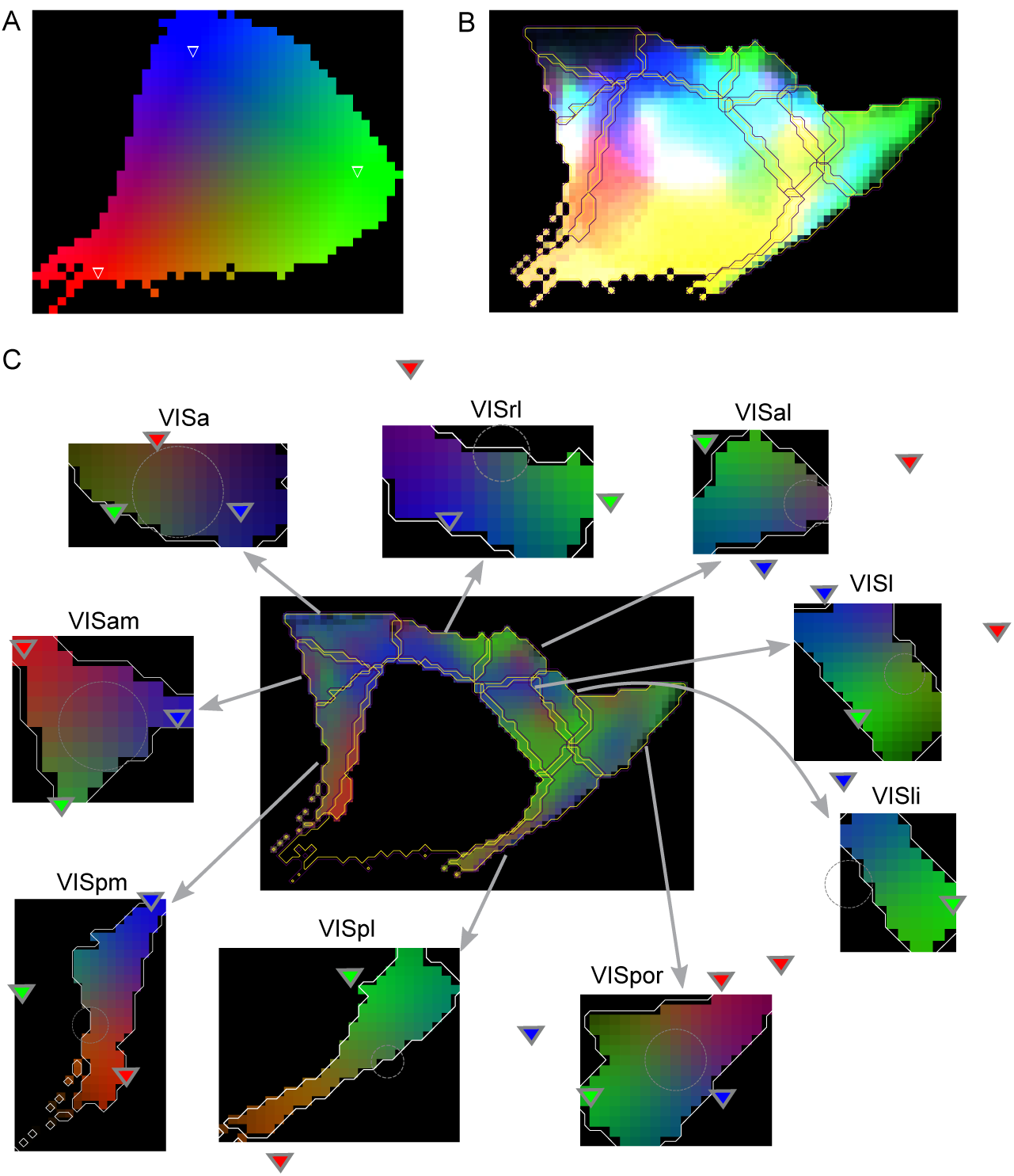
A. The primary visual area (VISp) and its defined source coordinate system. The three points defining the barycentric system are indicated as colored triangles. Each coordinate is associated with the indicated red, green or blue color channel to decide the color of each pixel in the region. B. The spatial structure of projections from VISp is indicated by coloring pixels in the surrounding regions according to the color in A of the area they are innervated from. C. Center: As in B, but the color of each pixel is normalized such that the sum of the red, green and blue channels is constant. Periphery: Target coordinate systems for the surrounding regions were fit to recreate the color scheme of the center, when colored as in A.

The results showed a clear non-random structure of targeting in the other regions (e.g. for projections from VISp: Fig. 5B). To parameterize this structure, we first normalized the color values of each pixel, dividing them by the total projection strength reaching that pixel from *src*. We set the denominator to a minimum of 25% of the maximum strength from *src* in the target region to ensure that weakly innervated parts of *tgt* would be depicted as such.

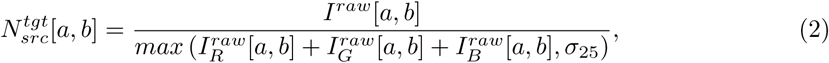

where *I*[*a, b*] denotes the pixel of image *I* at coordinates *a, b* and

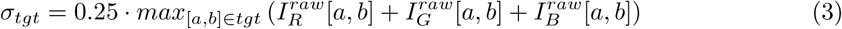

This represented a projection as pixels with normalized lightness, that faded to black in weakly innervated parts of the target region (Fig. 5C, center, 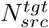). Next, we optimized a barycentric coordinate system in the 2d-projected target region to most closely recreate the color scheme observed in 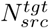 (Fig. 5C, periphery, 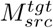). We then assume that a neuron at any coordinate in *C_src_* is mapped to neurons at the same coordinate in 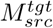. Thus, the two local coordinate systems, each parameterized by three points, together define the topographical mapping between regions *src* and *tgt*.

#### 3.3.2 Validating the mapping

We validated our predicted mapping against established data on the retinotopic mapping in the visual system. This is functional data on the mapping between a brain region and locations in the visual field instead of anatomical data on the projections between brain regions. Yet, we can use it for validation under the assumption that areas corresponding to the same location in the visual field are preferably projecting to each other.

Analyzing the retinotopy, Wang and Burkhalter (2007) found certain trends: In adjacent regions, points close to the boundary between them on both sides are mapped together. Also, a counter-clockwise cycle in one area is mapped to a clockwise cycle in an adjacent one. This change in chirality indicates that the mapping must contain a reflection operation. Juavinett et al. (2017) utilize this to identify borders between brain areas from intrinsic signal imaging of retinotopy. When we systematically examined the reflections and rotations in our predicted mapping (Fig. 6A), we found identical results.

**Figure 6:**
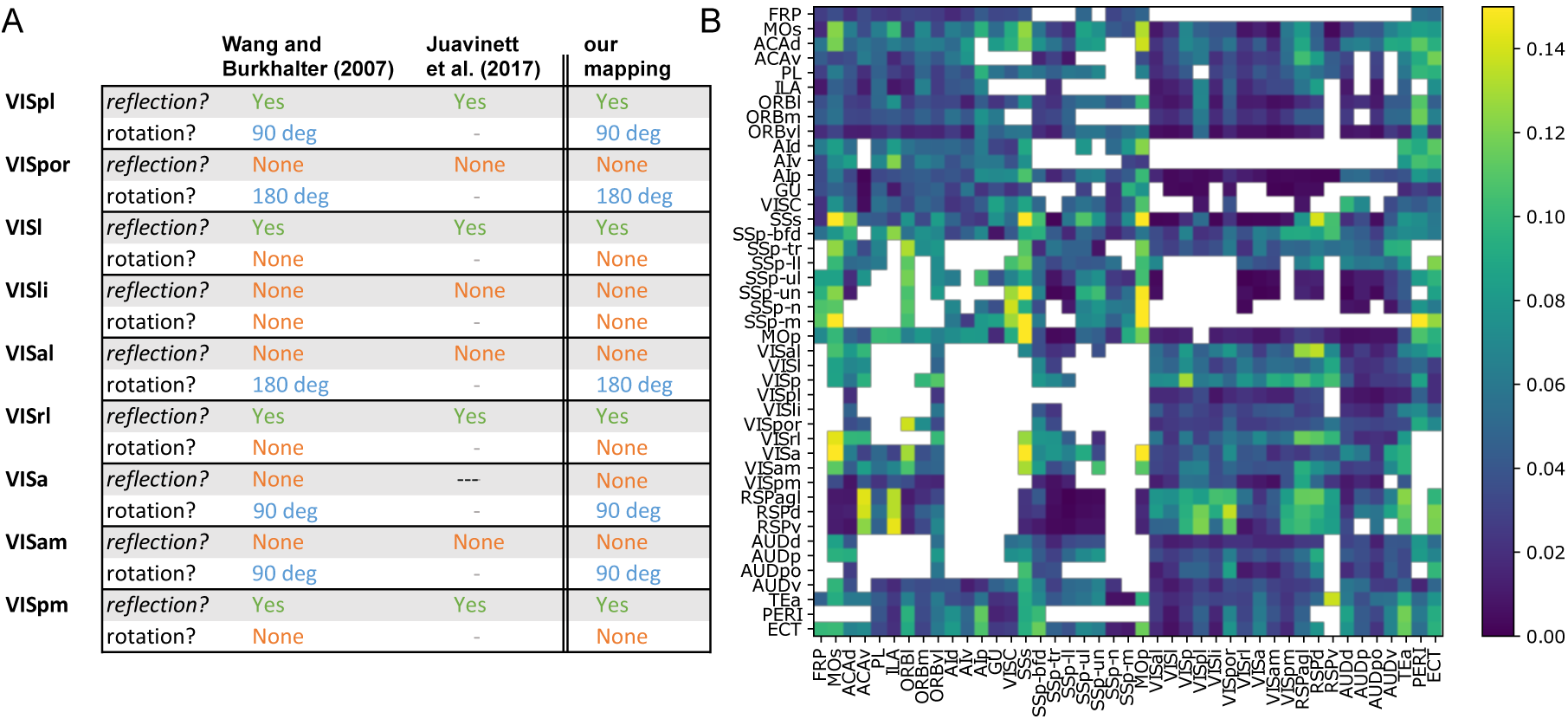
A: Comparing linear transformations from source to target coordinate system in our results to the ones of Wang and Burkhalter (2007) and Juavinett et al. (2017). B: Relative error of the mapping defined by the barycentric coordinate systems in the target area, compared to the data. Values along the main diagonal: for contra-lateral mapping; all others: ipsi-lateral mapping. Data shown where the sum of densities from all projection classes is above 0.025µm^−3^

Finally, we quantified to what degree barycentric coordinate systems in source and target region can capture the biological trends present in the projection data. As this type of mapping is always continuous and cannot capture non-linear trends such as exponential scaling, biological accuracy could be lost. To this end, we calculated the difference between the image of the target region, colored according to the target coordinate system, 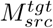, and the normalized image of the target region according to the projection data, 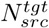. We defined the *relative error of a target coordinate system* as the sum of absolute differences of the two images, divided by their average and the number of pixels (Fig. 6B). We found that for over half of the projections the error was below 5% and the maximum error was 17%.

We observed that the largest error occurred for projections from and to the three largest regions: MOs, MOp and SSs, indicating that these regions could potentially be further broken up into subregions, similar to SSp (see Discussion).

#### 3.3.3 Parameterizing the width of the mapping

The mapping defined by the two local coordinate systems mapped points in one region to points in the other. However, due to the potentially large extent in the target region of single projection axons, the biological mapping is rather point-to-area than point-to-point. This results in a less strict mapping that the above method would depict as an image with slightly “washed-out” colors. For example consider a projection where 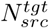 results in all grey pixels. This could either indicate a strict mapping where only the most central point of the source region (associated with grey) is mapped to the target region (i.e. the mapping incompletely covers the source region), or alternatively the whole source region is mapped to the target region, but the topographical structure of mapping is so weak that it becomes all but invisible.

Indeed, we found for most projections low saturation values in 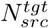 and consequently the optimal solution for the target coordinate system 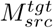 would place all three defining points outside the target region. This would lead to more peripheral points of the source region being mapped to locations outside the target region, i.e. to no target neurons at all. However, we assumed that low saturation values were rather a result of a large extent of projection axons leading to a weak mapping. We therefore added another objective to the optimization procedure for 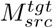: minimizing the fraction of the source region that is mapped to points outside the target region. To compensate, we defined points in the source region to be mapped to 2d-gaussian kernels at their target location instead of a single point. The width of the gaussian was optimized such that a convolution of 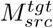 with the same gaussian resulted in the same distribution of saturation values as 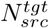.

#### 3.3.4 Incomplete mapping

To model the mapping between brain regions from the voxelized connectivity data, we had to make a strong assumption on the nature of said mapping: That it is continuous with only linear scaling. We also assumed that the mapping tends to cover most of the source region (see above), although the model will account for it if the data strongly indicates so. For example, feedback projections in the somatomotor system tended to cover the source regions only incompletely (Fig. 7 A, columns “data”). Note, how SSp-ll was only targeted by projections from neurons in MOp associated with red, and from neurons in SSs associated with red or green. Conversely, SSp-m were targeted by the regions of MOp and SSs associated with green or blue. This is not a surprise, as the primary regions are broken up into representations of individual body parts and the higher-order regions are not. The model captured this trend through target region coordinate systems where some of the defining points landed far outside the region (Fig. 7 A, columns “model”).

**Figure 7:**
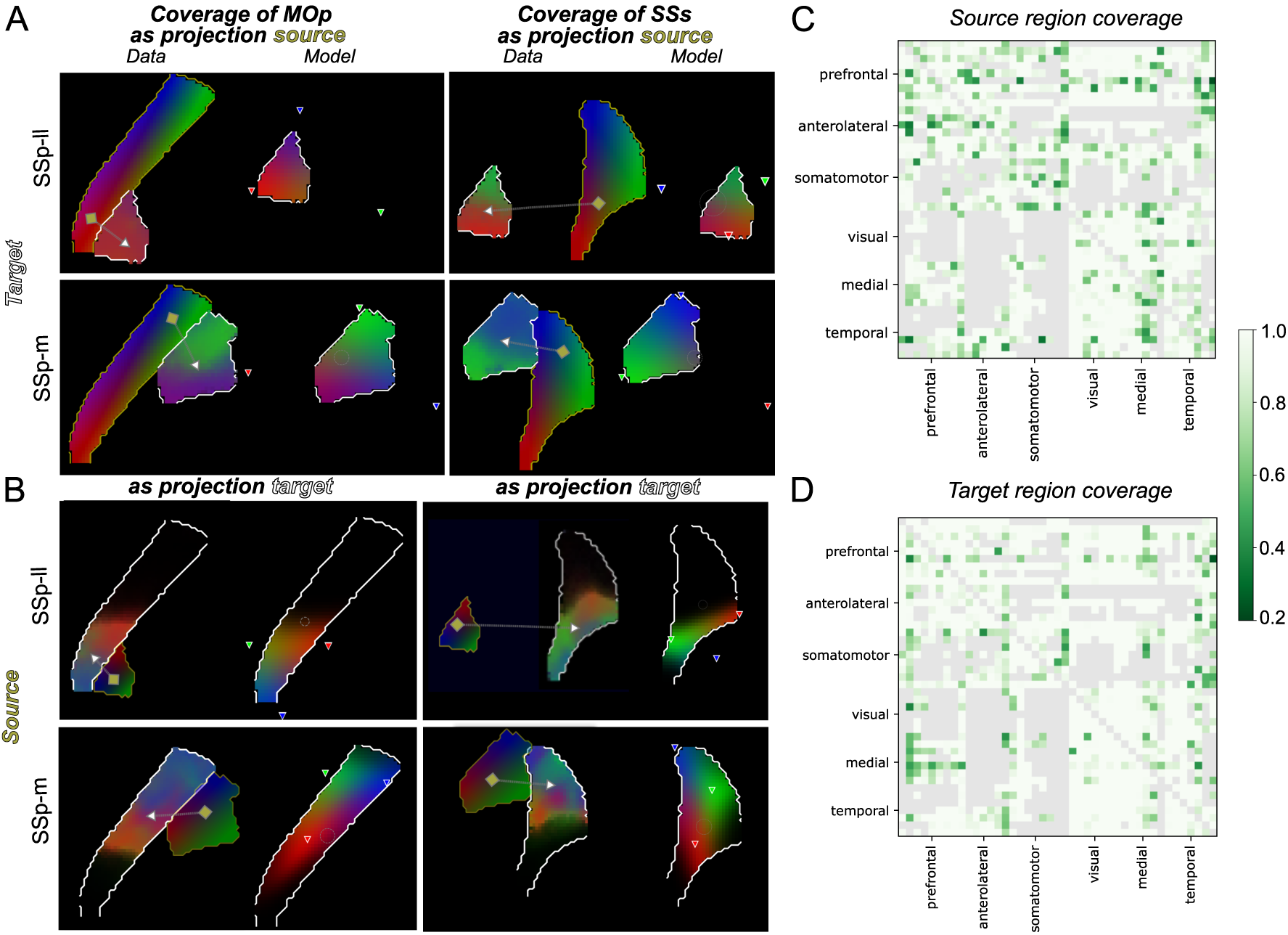
A. Exemplary feedback projections in the somatomotor module only incompletely cover the source regions. Projections are MOp to SSp-ll, SSs to SSp-ll, MOp to SSp-m and MOs to SSp-m (top left to bottom right). Relative projection strengths (left) and the resulting barycentric model (right), including the defining points (colored triangles) are depicted. B. Exemplary feedforward projections in the somatomotor module only incompletely cover the target regions. As in A, but source and target region are swapped (SSp-ll to MOp, SSp-ll to SSs, SSp-m to MOp, SSp-m to SSs). C. Fractions of the source regions covered by ipsi-lateral projections. Grey areas depict projections weaker than a threshold value or connectivity within a region. Ordering as in Fig. 6 and Table S1. D. As C, but coverage of the target region.

Conversely, we made no assumption of complete coverage of the target region. While it was largely complete in the upstream pathways of the visual system (Fig. 5), according to the data this was not the case for feedforward projections in the somatomotor system, and the model captured this trend (Fig. 7 B). Regions as targets of feedforward projections were only incompletely covered, and in fact the same parts of the regions were engaged in both feedforward and feedback projections with a given primary somatosensory region. Furthermore, as a group, the primary somatosensory regions engage all parts of other regions about equally. For example the parts engaged by SSp-ll and SSp-m are non-overlapping and together completely cover MOp and SSs (Fig. 7 A vs B).

Overall, mapping coverage was complete for 50% of projections on the source side and for 62% of projections on the target side. Over the whole neocortex, average source region coverage was 89% and target region coverage was 84% (Fig. 7 C, D). Incomplete target region coverage was taken into account in the scaling of anatomical projection strengths (see above).

### 3.4 Constraining projection types

Thus far, we have considered constraints on the spatial structure of projection synapses on a global scale (the macro-connectome matrix) and a local scale (the layer profiles and the mapping). The topographical mapping also limited which individual neurons in a target region can be reached by a given neuron in a source region, severely constraining the topology of the potential connectome graphs on a local scale. Yet, an important aspect of neocortical connectivity not yet considered is which combinations of *regions* are reached by single source neurons (Han et al., 2018). Even if we know which regions are innervated by the *population* of neurons in a given region, it does not mean that every single neuron that is part of the population innervates every single of those regions. Instead, each neuron innervates a subset of the connected regions and we call that subset its projection type or *p-type*. It is unclear to what degree the process is pre-determined or stochastic, and if it is stochastic, what mechanisms further shape and constrain the randomness. This is a complex problem, as a region such as SSp-tr innervates 27 other regions, yielding 2^27^ = 134217728 potential p-types.

To tackle this problem, we analyzed reconstructed axons made available by the MouseLight project at Janelia. These are whole-brain neuron reconstructions of cortical neurons that include their long range projections. We first classified their neuron types, then placed the axons in the context of the Allen Brain Atlas and finally evaluated the amount of axonal length projecting into the 43 ipsi- and 43 contralateral brain regions. We could now use this data to try and constrain a model of the targeting of individual axons.

Figure 8A shows an example of 61 analyzed axons originating in MOs. The scale of the p-type problem is clear at first glance: Only a single combination of innervated regions is repeated in this data set, all others represent unique p-types. Yet, a structure is also apparent: While only 11 out of the 61 axons innervate the visual or medial modules, the ones that do tend to innervate more than a single of their regions; up to 8 regions for axon number 9. This indicates a positive statistical interaction between the regions of those modules. Moreover, it appears that the projection strength (Fig 8A first row) is a strong predictor of the probability that any given axon innervates a region (*innervation probability*), indicating that a projection is strong because many neurons participate in it, not because of few participating neurons with strongly bifurcating axons in the target region.

**Figure 8:**
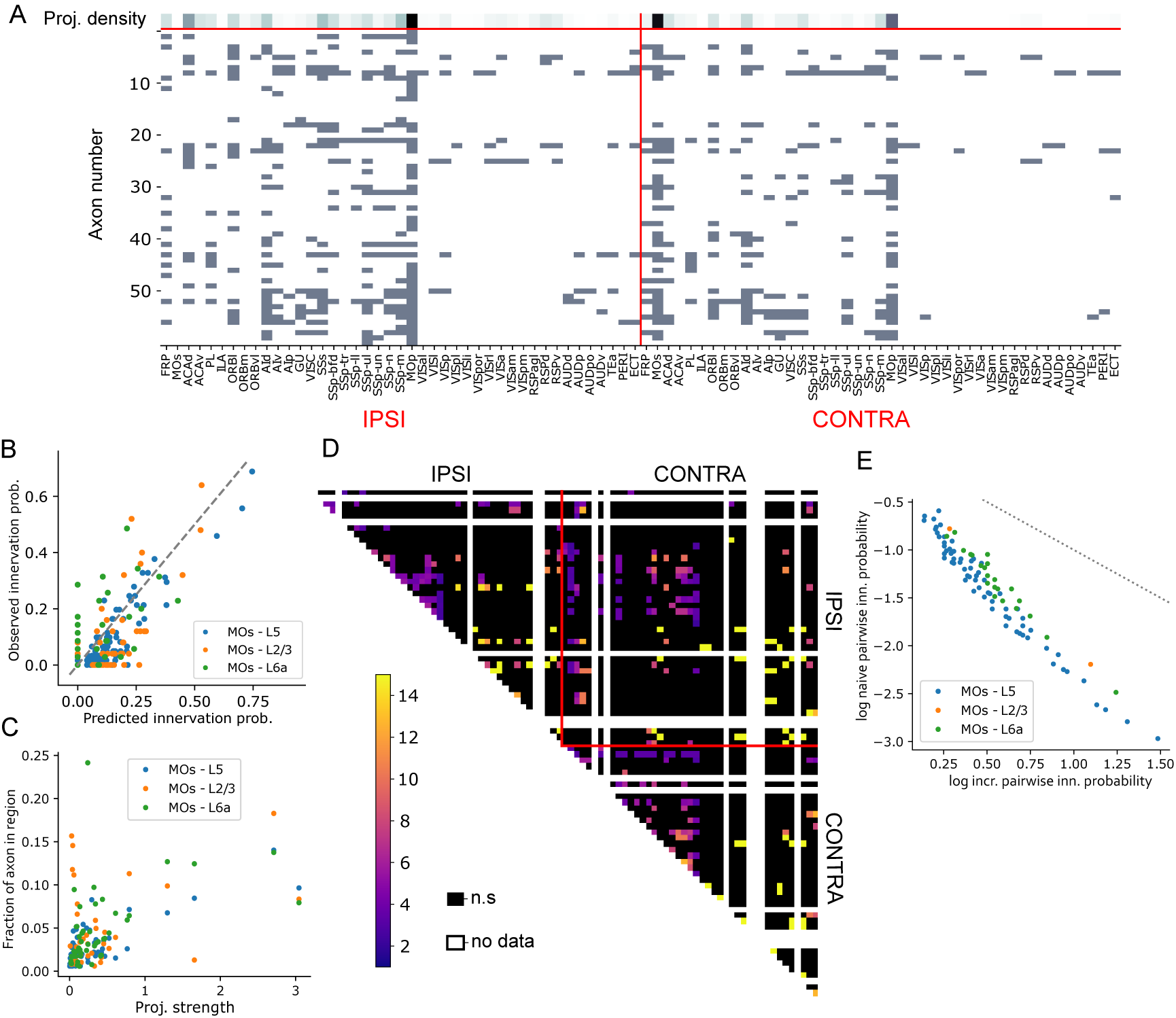
A. Projection density according to Harris et al., 2018 (top row) and brain regions innervated by 61 reconstructed axons (rows) indicated by gray squares. B. Probability to innervate individual brain regions, predicted from the normalized projection strength as proportional to the square root of the normalized projection strength, against the observed innervation probability. C. Normalized projection strength against the mean total length of axon branches in individual brain regions. D. Observed interactions between the innervation of individual brain regions, i.e. increase in innervation probability of one region when the other is known to be innervated. E. Increase in innervation probability as in D against the innervation probability of a pair of regions under the assumption of independence. Gray dotted line indicates the point where the product of independent probability and increase is 1 that can logically not be exceeded. All innervations and projection strengths in this figure are for projections from MOs.

Next, we analyzed these observations systematically. We only had for the source region MOs a sufficient number of reconstructed axons to robustly estimate the innervation probabilities. When we considered the relation between the strength of a projection from MOs and the probability that a neuron in MOs participates in it, we found that the best predictor was the *normalized projection strength*, i.e. the amount of signal (axon) in the target region normalized by the volume of the source region. Specifically, the predictor of innervation probability was 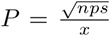, where *nps* refers to the normalized projection strength and *x* to a projection class-specific factor determined by a linear fit (*x* = 2 for projections from L2/3 and L6, 3 for projections from L4 and L5PT and 4.5 for projections from L5IT; Figure 8B, *p* = 3⋅10^*−*9^, pearsonr). Projection strength being a predictor of innervation probability is in line with the findings of Han et al. (2018).

Conversely, the projection strength was less a predictor of the axon length in a target region for individual axons innervating the region (Figure 8C). While there was a significant correlation (*p* = 1.5 · 10^*−*3^, pearsonr), the strength of the effect was much weaker than for predicting the innervation probability. Particularly, while a very weak projection strength predicted low innervation probabilities, this correlation broke down quickly for innervation probabilities above 3%. Taken together, this means that the strength of a projection is mainly a result of more neurons participating in it and less of the length of individual axons in the target region. But most importantly, assuming the principle holds for other brain regions as well, we were able to predict the first order innervation probabilities for all combinations of source and target region.

Next, we analyzed statistical interactions of the innervation probabilities for axons originating in MOs. For pairs of target regions (42 ipsi- and 43 contra-lateral), we evaluated the null hypothesis that their innervations are statistically independent, and if it was rejected (*p* ≥ 0.05) calculated the strength of the statistical interaction as the conditional increase in innervation probability 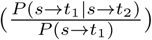. We found significant interactions for 283 pairs (Figure 8D), with some strengths exceeding a 15-fold increase. However, there were several problems preventing us from simply using these observed interactions to constrain connectivity. First, we only had data for axons originating from one of 43 brain regions and it is likely that interactions differ for source regions. Second, the data was incomplete, as some target regions were not innervated by a single reconstructed axon (Figure 8D, white patches) and others were based on only a single or two axons. Third, evaluating 86 ⋅ (86 − 1)*/*2 = 3655 potential interactions based on only 61 data points (i.e. axons) is statistically inherently unstable and likely to dramatically overfit.

#### 3.4.1 A model to generate “p-types”

Instead, we tried to use the available axon data to develop a conceptual model of how the interactions arise. We first observed that the largest interactions strengths occurred for target regions in the medial and visual modules that are otherwise only weakly innervated. Evaluating this observation systematically, we found that indeed the strength of an interaction was strongly negatively correlated with the product of the first order innervation probabilities of the pair (Figure 8E). Second, we observed only conditional increases in innervation probability (values *≥* 1), i.e. innervation of pairs of brain regions is not mutually exclusive.

One model explaining both our observations is the following: Consider a tree with the brain regions in both hemispheres as the leaves. Let each edge in the tree be associated with a probability that the edge is successfully crossed by an axon, these probabilities can be different in both directions of the edge. To generate the set of innervated regions for a random axon, start at the leaf representing its source region and then consecutively spread to other nodes further into the tree along its edges with the probabilities associated with the edges. Once it has been decided that an edge is *not* crossed, it cannot be crossed in future steps. Every leaf reached this way is then considered to be innervated by the axon.

If we set the length of an edge in this model to the negative logarithm of the associated probability, then the first order probability that a region *T* is innervated by an axon originating in region *S* is easily calculated:

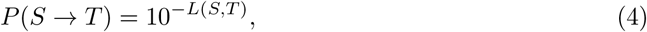

Where *L*(*S, T*) denotes the length of the shortest path between *S* and *T*.

Similarly, the increase in conditional innervation probability of *T*_1_ and *T*_2_ is given as:

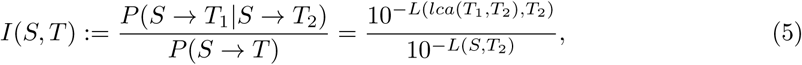

Where *lca*(*T*_1_*, T* 2) is the lowest common ancestor of *T*_1_ and *T*_2_.

Due to the underlying tree structure, the lowest common ancestor is always an inner node that is closer or of equal distance to *T*_2_, therefore the strengths of interactions are always larger than one indicating an increase of innervation probability, which is in line with our earlier observations.

#### 3.4.2 Fitting the p-type model

Fitting the model consisted of two steps: First, we generated the topology of the tree using the *normalized connection density* of projections, i.e. the amount of signal (axon) in the target region normalized by the volume of both source and target region. Specifically, we used the Louvain heuristics (Rubinov and Sporns, 2010) with successively decreasing values for the *gamma* parameter to detect successively larger communities in the matrix of normalized connection densities (see Methods). Then we optimized the probabilities associated with edges using the first order innervation probabilities predicted from the *normalized connection strength* of projections as in Fig. 8B. These predictions then served as constraints on the path lengths between leaves. Specifically, we locally optimized the edges in small motifs consisting of two sibling nodes and their parent, based on differences in the distances of the siblings to all leaves (see Methods). Note that the data to fit the model was the dataset of Harris et al. (2018) and the reconstructed axons of the Mouselight dataset were only used to inspire the model, to fit the method to predict first order innervation probabilities, and for subsequent validation. Figure 9B shows the resulting trees for all projection classes along with the predicted statistical interactions.

**Figure 9:**
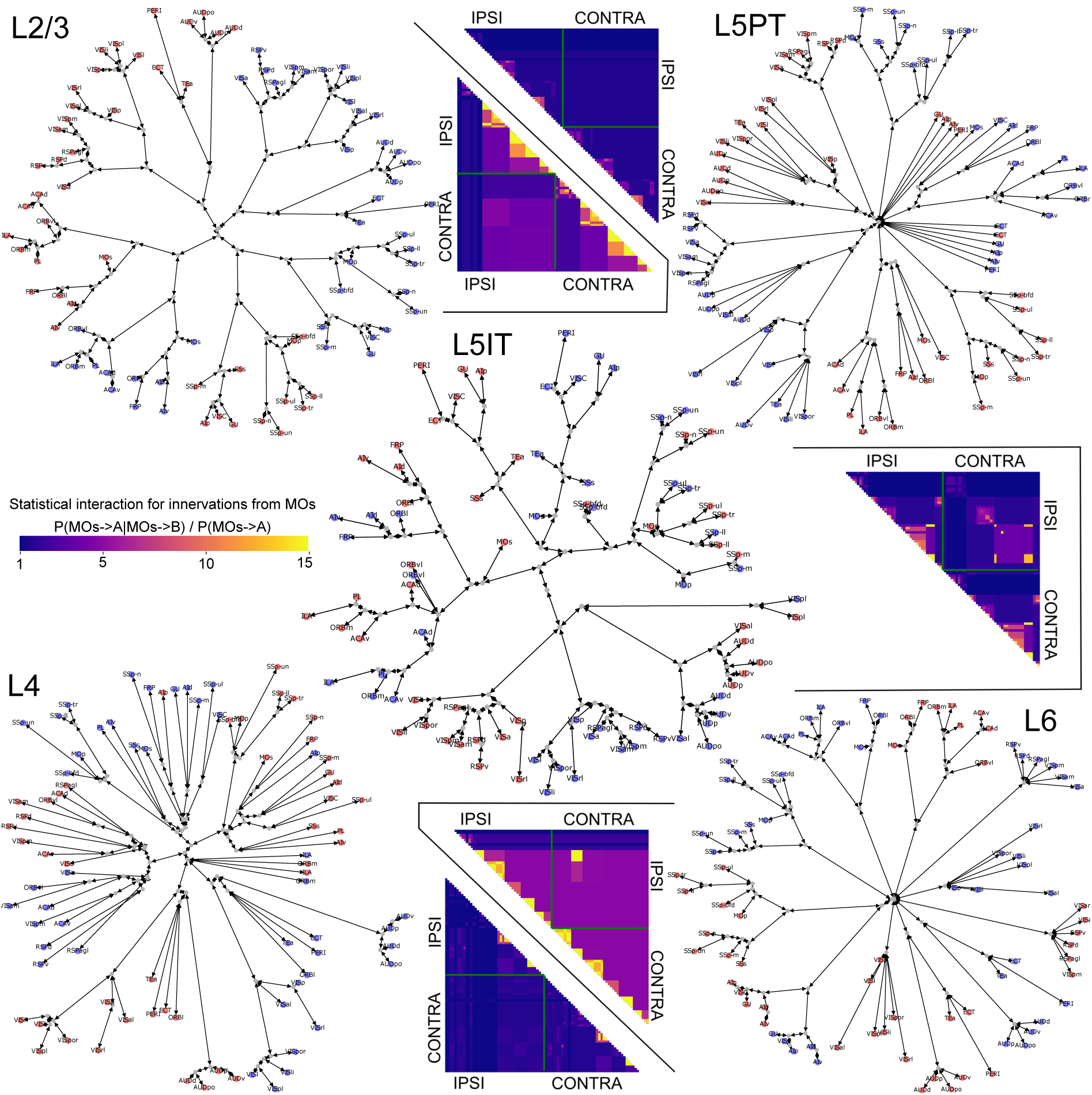
The final tree models for all projection classes. Edge lengths depicted are the mean of lengths in either direction. Blue / red nodes are leaves representing brain regions in either hemisphere. Depicted alongside each tree are the predicted statistical interactions between innervations of ipsi- and contra-lateral regions from MOs as in Fig. 8D.

#### 3.4.3 Validating the p-type model

We used the fitted model to generate 10000 profiles of brain region innervation for axons originating from L5 of MOs and MOp. Figure 10A compares a number of randomly picked profiles against the data from reconstructed axons for both regions. As the model was constrained with the predicted first order innervation probabilities, it manages to recreate the observed high-level trends: Strong innervation of the ipsi- and contralateral prefrontal, anterolateral and somatomotor modules; weaker, but highly correlated innervation of the other modules.

**Figure 10:**
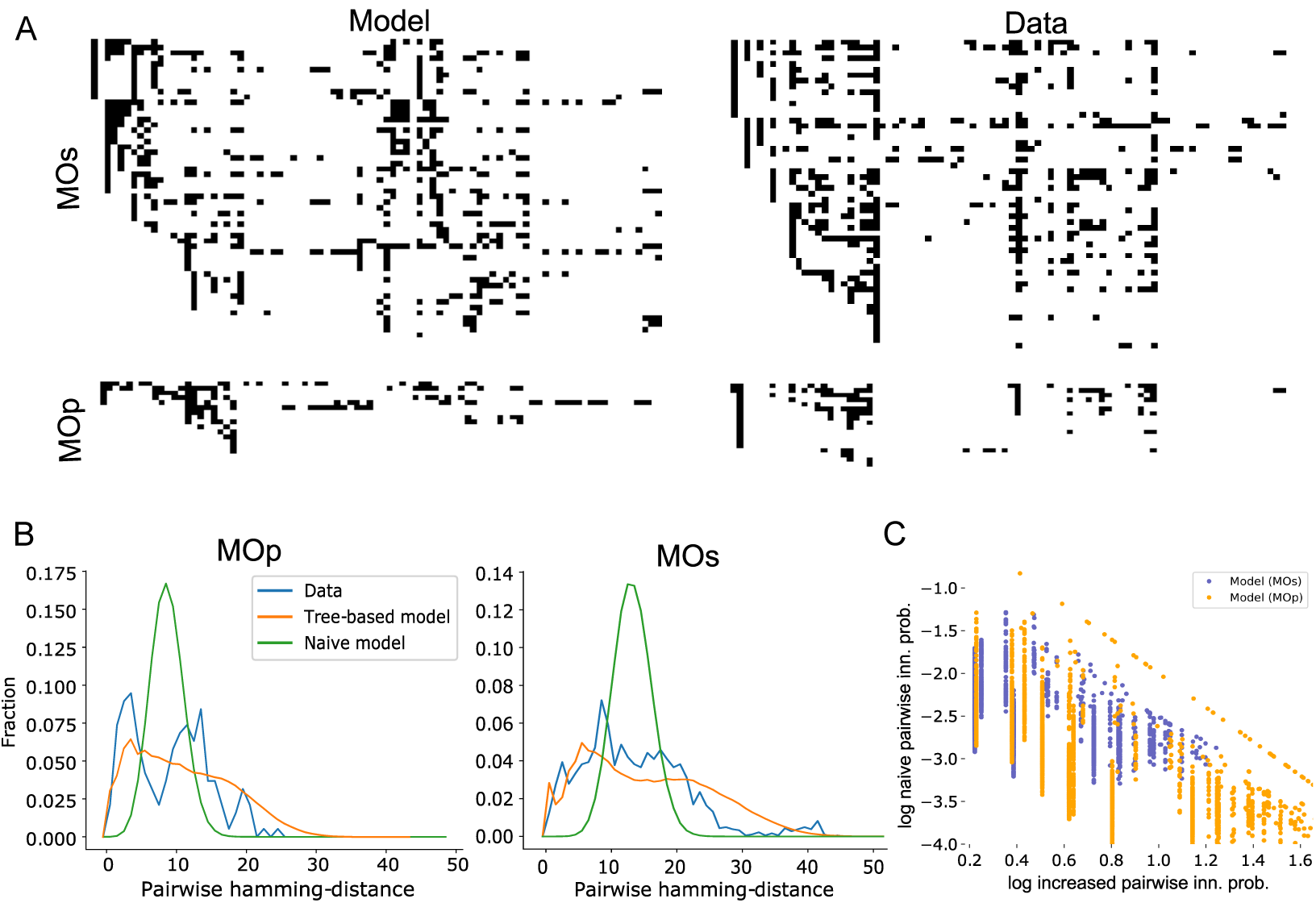
A. Examples of innervation of brain regions predicted by the model (left column) and of reconstructed axons (right column). Sampled axons along the y-axis, brain regions along the x-axis. A black pixel indicates that an axon is innervating the indicated region. Top row: axons originating from L5 of MOs, bottom row: from MOp. B. Pairwise distances (hamming distance) between the profiles of brain region innervation. Blue: Data from reconstructed axons (see A); orange: from 10000 profiles sampled from the tree-based model; gree: from 1000 profiles sampled from a naive model taking only the first order innervation probabilities into account. Left: for axons originating from MOp; right: from MOs. C. Increase in innervation probability against the basic innervation probability as in Figure 8E.

To test the model further, we calculated the pairwise hamming distances between innervation profiles from reconstructed axons and from the model (Figure 10B). We also compared the data against a naive model using only the observed first order innervation probabilities and assuming no interactions. We found that the naive model resulted in a narrow, symmetrical distribution with a single peak at around 9 (MOp) or 13 (MOs). In contrast, the axon data led to a much wider, asymmetrical and long-tailed distribution that was much better approximated by the tree-based model. The difference between the distribution resulting from the tree-based model and the axon data was, in fact, not statistically significant (MOp: p=0.44; MOs: p=0.12; kstest).

Using the tree-model, we could predict the strengths of interactions as described in equation 4 (Figure 9). When comparing the strength of the interactions against the naive innervation probabilities without interactions, we found in the model the strong negative correlation that was present in the axon data (Figures 10C, 8E). For the model we found more data points towards the lower left corner of the plot that indicates low naive probability and low increase. The lack of such points in the data from axon reconstructions can be explained by the fact that points associated with extremely low probabilities are unlikely to show up in a relatively small sample of reconstructed axons.

As a final validation, we compared the model against the results of Han et al. (2018), which considered brain region targeting of single axons originating from VISp. We have not taken into accounts axons from this source region when we formulated or fitted the model, making this a powerful validation of the generalization power of the model (Fig 11). Comparing the number of visual regions innervated (out of VISli, VISl, VISal, VISpm, VISam, VISrl) by individual axons originating in layer 2/3 of VISp, we find comparable results (Fig 11 A). Although in the model the mean number of regions innervated is slightly higher (1.84 vs 1.7 (fluorescence-based) or 1.56 (MAPseq)) we find the same roughly binomial distribution where fractions decrease with increasing number of innervated regions.

**Figure 11:**
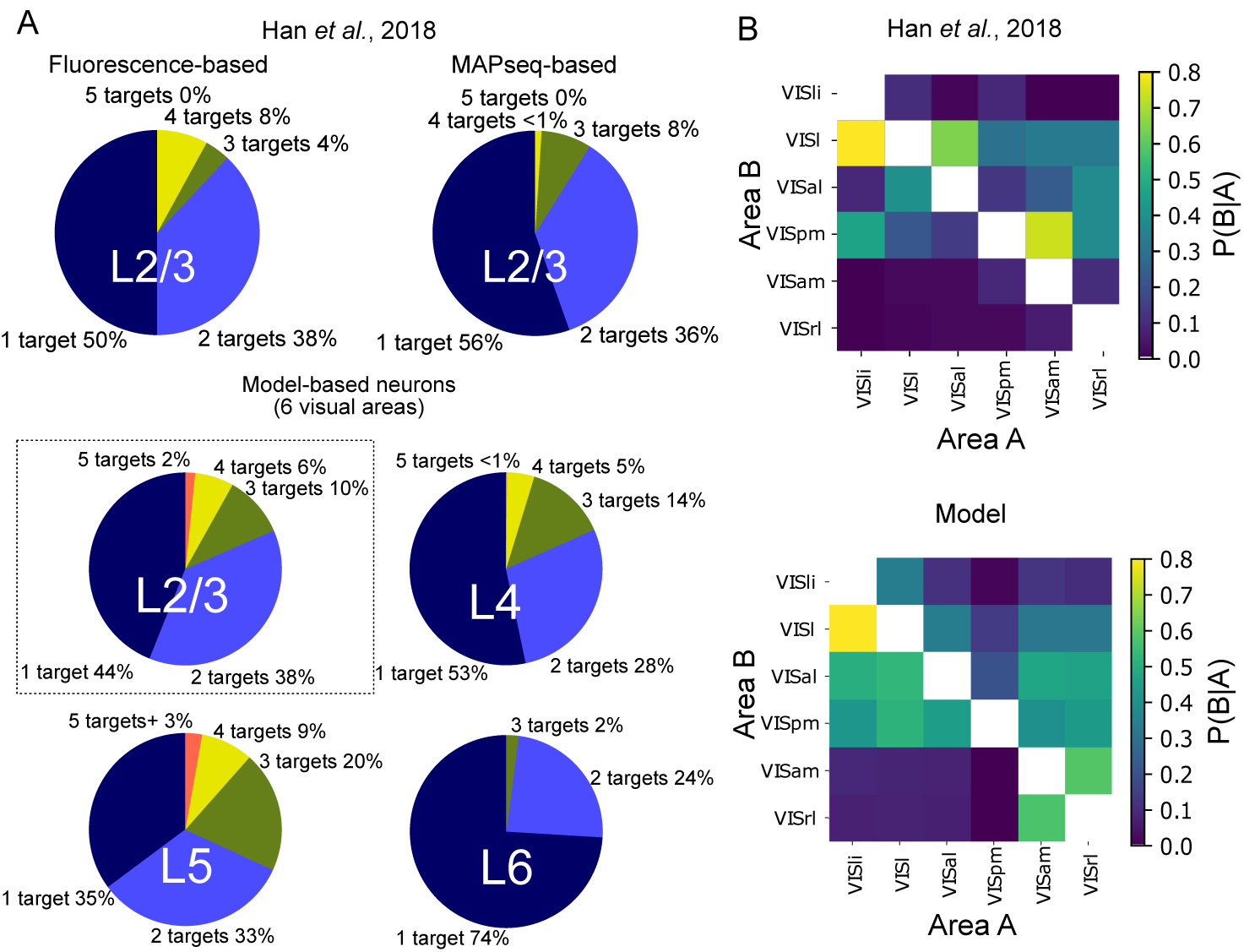
Validation of the tree-model against the results of Han et al. (2018). A: Top, results of Han et al. (2018) in terms of the number of visual areas innervated by single axons originating in layer 2/3 of VISp. Bottom, corresponding results of the tree-model for axons originating in layers 2/3, 4, 5 and 6 (top left to bottom right). B: Top, results of Han et al. (2018) in terms of common innervation of pairs of visual brain areas by axons originating in layer 2/3 of VISp. Bottom, corresponding results of the tree-model.

We were also able to predict this distribution for axons from other layers using our model. We predict similar shapes of the distribution with an even higher mean for layer 5 and a lower mean for layer 4 and especially layer 6. Next, we also considered the statistical interactions between the six visual target regions (Fig 11 B). Again, we found overall comparable conditional probabilities, with a comparable structure, although strong common innervations of regions VISl and VISal and VISpm and VISam were underestimated.

Taken together, the tree-based model to generate p-types has generalization power for other brain regions and can serve as a suitable null model of the region targeting of single axons.

### 3.5 Connectome consistency and availability

We had scaled the strengths of projections to obtain a target value of 88.74 billion local and long-range synapses (see methods). Our established algorithm to derive local connectivity within a brain region (Reimann et al., 2015) resulted in 23.05 billion synapses, predicting a ratio of 23.05 billion to 65.69 billion 26≈ to 74% of local to long range connectivity. This is roughly in line with our earlier estimates of approximately 80% long range connectivity into non-barrel somatosensory cortex, based on total spine densities (Markram et al., 2015) and based on structural excitatory to inhibitory balance of individual neurons (Reimann et al., 2017). Another way to estimate the fraction of local vs long-range inputs is to directly consider the projection matrices of Harris et al. (2018). The matrices contain entries for the synapse densities of connectivity within a region along the main diagonal, that we compared to the sum of all other entries for any target region. The resulting estimates of local vs. long-range connectivity ranged from 5% - 95% to 35% - 65% between regions with an average of 19% - 81% (mean across target brain regions, weighted by region volume), in line with our predictions.

We parameterized all constraints listed above - projections strengths, layer profiles, topographical mapping and p-types - for all projections and exported the results as a *long-range projection recipe* in the machine-readable yaml format. This recipe is publicly available under https://portal.bluebrain.epfl.ch/resources/models/mouse-projections.

### 3.6 Connectome instantiations

Finally, we developed a stochastic algorithm to generate instances of a neuron-to-neuron connectome that fulfills all constraints in the long-range projection recipe and used it to connect a model of the entire mouse neocortex. Briefly, it generates for each projection a voxelized volume of target synapse densities, based on layer profiles and projection strength. Next, it places synapses on randomly picked dendritic segments in each voxel until the target density is reached. Finally, it connects each placed synapse to a presynaptic neuron based on the topographical mapping and p-types (for details see Methods). We considered slender tufted and untufted pyramidal cells in layer 5 to participate in projection class L5IT and half of the thick tufted layer 5 pyramidal cells in L5PT, with the other half participating in L5CT, which is not covered by the present, purely cortical model. Pyramidal cells in other layer all participated in the corresponding projection class.

As a result, we obtained for each connectome instance 88 billion modeled synapses, for each of which we know the presynaptic neuron, postsynaptic neuron and the exact location on the postsynaptic morphology. Consequently, we were able to perform for example retrograde tracer experiments revealing complete maps of the neocortical neurons innervating small brain volumes (Fig. 12). The *in-silico* method allowed us to target volumes several orders of magnitude smaller than possible with experimental methods (Gămănuţ et al., 2018), down to the innervation of individual neurons with sub-cellular resolution (Fig. 13). In the future, this will allow us to simulate the electrical activity of individual neurons, entire regions or of the entire neocortex. We made sparse connection matrices of seven instances of the predicted null model of neocortical long-range connectivity publicly available under https://portal.bluebrain.epfl.ch/resources/models/mouse-projections.

**Figure 12:**
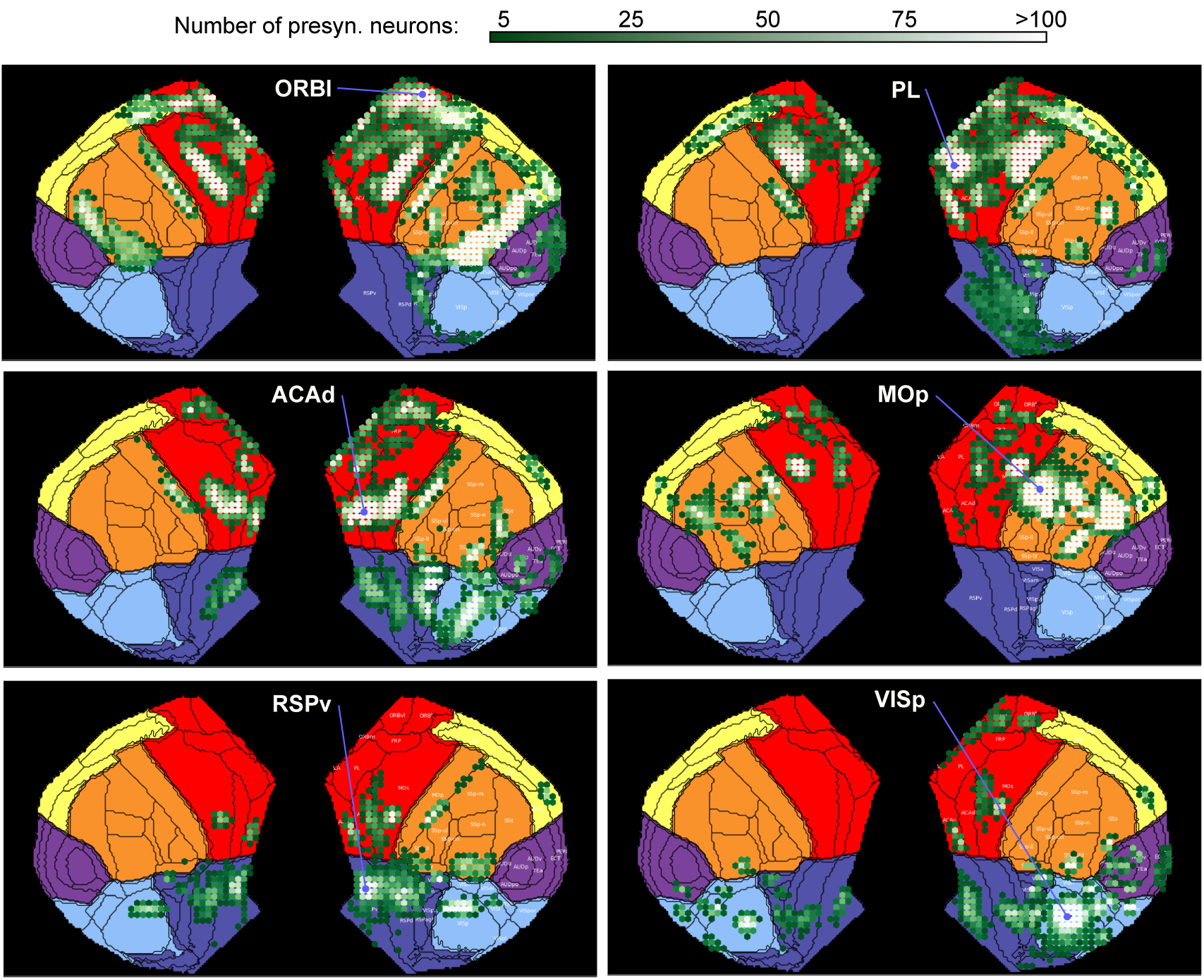
In-silico retrograde tracer experiments. Locations and numbers of neurons innervating around 100 neurons in a small volume (1.8 ⋅ 10^*−*4^*mm*^3^) at the indicated locations in various representative brain regions.

**Figure 13:**
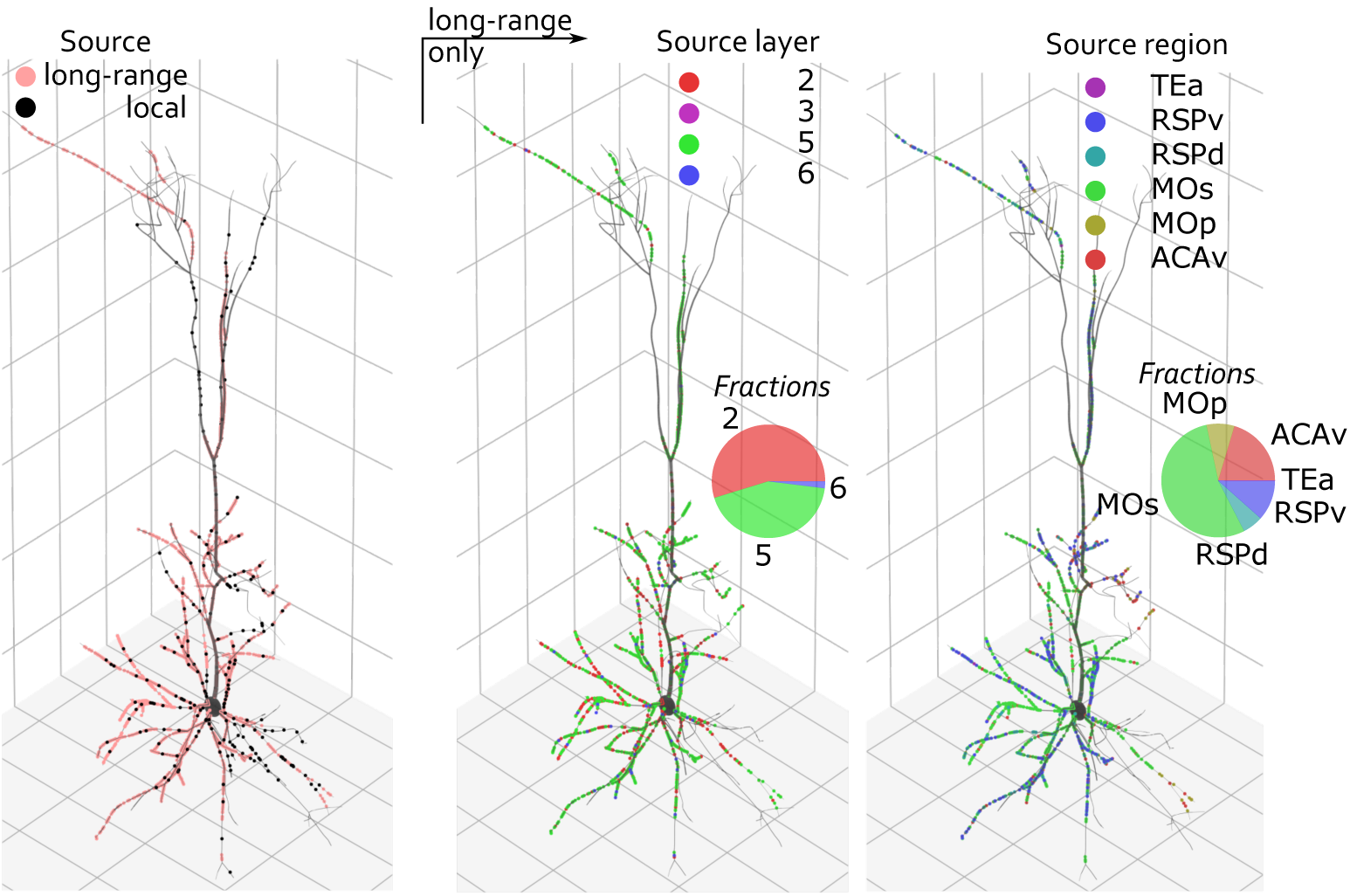
Modeled synaptic innervation of an exemplary PC in layer 5 of ACAd in terms of the origin of each synaptic contact. Local vs. long-range (left), by source layer (middle), or by source region (right).

## Discussion

We have developed a way to generate statistical instances of a whole-neocortex mouse micro-connectome. This approach takes into account the current state of knowledge on region-to-region connectivity strengths, the laminar pattern of projection synapses, the structure of topographical mapping between regions and the logic of regional targeting of individual projection axons as derived from over 100 whole-brain axon reconstructions, and a comprehensive meso-scale model of projections, built from thousands of experiments (Harris et al., 2018). Combining these data with a morphologically detailed model of neocortex (Erö et al., 2018), has allowed us to statistically predict connections with sub-cellular resolution, i.e. including the the locations of individual synapses on dendritic trees. Our approach is timely, as it leverages and integrates three very recent, publicly available datasets. Furthermore, its flexibility and modularity will allow it to readily use future datasets in place and in addition to the currently used ones. The resulting wiring diagram allows fundamental questions to be addressed, such as the nature and dynamics of clinically relevant brain rhythms as well as hierarchical interactions in cortex, which are fundamental for understanding cortical coding and whole-brain regional dynamics.

As the available data on this topic remains sparse, our approach was as follows: We assumed the formation of the connectivity is a stochastic process selecting one out of a space of possible wiring diagrams, and then seeked out biological principles and rules that consecutively restrict this *space of biologically viable wiring diagrams*.

The principles we identified were based on biological data, but also a number of assumptions. The assumptions were necessary to break down the scale of the problem, to interpret the data (data assumptions) and structure it into principles (structuring assumptions), to formulate principles mathematically (modelling assumption), and to apply them to infer missing data (generalizing assumption). In order to interpret the resulting micro-connectome and predictions, one needs to first understand these assumptions, summarized below:

- **Data assumption**: Connectivity is symmetrical between hemispheres, both within and across hemispheres.
- **Data assumption**: The amount of fluorescent signal is directly proportional to axon length in a region and the density of synapses per unit axon length is uniform across all neocortex. That is, the density of synapses per *µm*^3^ is a constant multiple of fluorescent signal per *µm*^3^
- **Data assumption**: Connection matrices for individual projection classes are version of the wild type matrix, where submatrices are scaled by individual values, and that sum up to the wild type matrix. This assumption could be removed with more complete data on the projection classes.
- **Data assumption, generalizing**: First-order innervation probabilities for single axons can be predicted from the projection strength of the whole population in the source region. Validated in motor regions, we generalize the principle to all source regions.
- **Structuring assumption**: The five projection classes considered are suitable to describe and parameterize long-range connectivity.
- **Structuring assumption**: Each projection follows one of the six “prototype” layer profiles.
- **Structuring assumption**: A clear distinction between local connectivity within a region and long-range connectivity across region, where local connectivity follows the principles outlined in (Reimann et al., 2015) and long-range connectivity the principles outlined here.
- **Structuring assumption**: The brain parcellation scheme of the Allen Common Coordinate Framework is suitable to describe and parameterize the long-range connectivity.
- **Modelling assumption**: The mapping of connections between regions is generalized topographical, continuous, and any scaling is linear. However there is no assumption that it completely cover the source or target region. Note that we have quantified the amount of detail lost due to this.
- **Modelling assumption, generalizing**: The targeting of brain regions by individual axons can be explained by a tree-based model, generated from projection strength matrices. Validated for source regions VISp, MOs and MOp, we generalize to all other regions.
- **Inherited assumptions**: Assumptions that went into the construction of the voxelized connectivity model.
- **Inherited assumptions**: Assumptions that went into the construction of the morphologically detailed microcircuit model of Markram et al. (2015).
- **Implicit assumption**: The constraints on long-range connectivity we considered are complete, that is no other biological principles restrict the space of viable wiring diagrams. This is almost certainly not true.

As more and more data becomes available, fewer data assumption will be required. For example, we had to assume symmetry to build a full neocortical model because the chosen biological dataset was focused on the right hemisphere. With the addition of more data points in the left hemisphere, the assumption could be abolished and potential asymmetries revealed. Indeed, there is evidence of lateral asymmetries, also in non-human animals (Corballis, 2009), although it is unclear to what degree they are population-level tendencies that can be captured by a model based on data pooled from many individuals.

Structuring assumptions were made to structure the data, and reduce its complexity or to break up the problem to reduce its scale. For example, the clear split between local and global connectivity allowed us to tackle the individual sub-problems separately. These are shortcuts that are not inherently required and could potentially be phased out in future, improved versions of this model. The first candidate for such a future improvement is derivation of layer profiles of projections. The current approach proved challenging, due to the great variability in the biological data as reported by the voxelized connectome model, and consequently remains imperfect. A supplementary approach would be to analyze individual whole-brain axons, similar to our approach to region targeting. However, this comes with the caveat that analyzing individual axons in the context of an average brain parcellation scheme - such as the Allen common coordinate framework - leads to inaccuracies. While this is unlikely to upset the results for region targeting, the more fine-grained layer targeting will be more affected. Finally, for a complete model, axons originating from many different brain regions and layers would be required. Ultimately, whole-brain electron microscopy holds promise to solve these issues.

Modelling assumptions are integral to the model and relate to hypotheses about the biological system. We consider an assumption generalizing if it has been validated for part of the neocortex and is used for all of it. We also inherit all assumptions that were made in order to generate the voxelized connectivity model that we used.

Finally, there is the implicit assumption of completeness, that our model captures all pertinent biological principles. We make no claim that this is true. This assumption is formally necessary for us to achieve the following modelling goal: **Given the assumptions, find the most general model that completely describes the data**. This way, the model will serve as a valuable null model to evaluate future findings against. For any data point invalidating the model, we will be able to pinpoint which assumption it violates and thus provide it context. To start this process, we made the model and data available to the public. The parameterized constraints on projection strength, mapping, layer profiles and individual axon targeting (i.e. the projection recipe), as well as stochastic instantiations of whole-neocortex micro-connectomes can be found under https://portal.bluebrain.epfl.ch/resources/models/mouse-projections.

We can already hypothesize about additional principles that might have to be added in the future. In terms of “targeting” of connectivity, we have implemented many aspects of spatial targeting of brain regions and locations within a region, but within those constraints it remains random. It is possible that similar rules apply for the incoming long-range projections, i.e. which set of brain regions individual neurons are innervated by, and possible interactions between incoming and outgoing. In that case, we will be able to extent our definition of *p-types* to be the concatenation of *incoming p-types* and *outgoing p-types*.

Further, it is possible that there is even more fine-grained logic in the targeting, expressed in the topology of neuron-to-neuron wiring across regions. For example, a bias for reciprocal connectivity that has been demonstrated for local connectivity (Song et al., 2005; Perin et al., 2011), could also exist for pairs of neurons in different regions.

In terms of the large-scale inter-area connectivity trends, i.e. the macro-connectome, our approach does not make any predictions, but is instead explicitly recreating the input data used. In principle, other data sources than Harris et al. (2018) could be used for this purpose. For example, Gămănuţ et al. (2018) report a cortical mouse macro-connectome that recreates biological trends, such as a lognormal distribution over several orders of magnitude of projection strengths. They argue that their data captures several projections that are missed by Oh et al. (2014) (and consequently also potentially by Harris et al. (2018), which is based on similar computational methods). As their data provides potential sub-area resolution (see their Figure S2), it could be used to also constrain the mapping and consequently serve as the basis of a stochastic micro-connectome predicted with our method, albeit without distinction of projection classes.

We implemented the targeting of individual brain regions with a tree-based model that conceptualizes the axon growing throughout the brain. While we do not make any claims about an anatomical basis of this conceptual model at this point, it may prove fruitful to relate it to anatomical fiber tracts in the future. It is possible that the fiber tracts implement a super-graph that the trees we identified for individual projection types are subgraphs of. More general projection classes may exist with a molecular encoding of which edges of the super-graph are present or absent. Indeed, genetic signatures of individual neurons have already been systematically analyzed and linked to axonal projections (Tasic et al., 2016).

The assumption of a continuous, linear mapping between regions appears to solidly recreate the projection data, with only three regions leading to significant error (Fig. 6; MOs, MOp and SSs). One explanation for the error would be that these regions contain subregions that each send and receive their own, continuous projections. Indeed, for the projections from SSp-ll and SSp-ul to SSs (Fig. 7B, right), we see several peaks of the green and blue color channels in the data, whereas a single continuous mapping can only generate single peaks. This is not surprising, as MOs, MOp and SSs are not broken up by body part, unlike SSp that it strongly interacts with. In the future, the projection data could thus be used to further break up these regions, at least for the purpose of analyzing projections. With more advanced analyses and more data it may even become possible to hypothesize a brain parcellation scheme *ab initio* based on projection data.

Even with the imperfections outlined above, the present model will lead to advances in our understanding of brain function, when employed in simulations of whole-neocortex activity. The explicit parameterization of the constraints will allow us to change parameters to assess their impact. For example, it is at this point unclear whether the targeting rules for individual axons (p-types) will have an effect on high-level brain activity. Similarly, we can investigate to what degree the relatively simple topographical mapping in the model is sufficient for the upstream propagation of spatial information from VISp. Steps into that direction can be undertaken both in morphologically detailed models and point neuron models using the publicly available model connectome.

## 5 Acknowledgements

We thank Lida Kanari for help with the validation of projection types, Joseph Knox for assistance with the projection from the Allen common coordinate framework into a 2d plane and Dimitri Rodarie & Max Nolte for feedback on the manuscript.

## 5.1 Author contributions

MWR developed, implemented and validated the techniques to constrain and parameterize the long-range micro-connectome, and to express the constraints in a machine-readable format. EM contributed to the development of these techniques. MWR generated the figures, and wrote the manuscript. MG developed and implemented the techniques to generate micro-connectome instances in a neocortex model based on the constraints. MWR and MG validated the microconnectome instances. YS and HL analyzed the axonal targeting of individual neurons from the Janelia MouseLight dataset. EM and HM provided guidance and contributed to writing the manuscript.

## 6 Methods

Unless noted otherwise, data from the voxelized mouse connectivity model of the Allen Institute was accessed using the “mcmodels” python package provided by the authors (https://github.com/AllenInstitute/mouseconnectivitymodels.git).

### 6.1 Volumetric synapse densities of projections

We formulated a target mean density of synapses of 0.72*µm^−^*^3^ in the model, as measured by Schüz and Palm (1989). Multiplied with the neocortex volume of the isocortex in the Allen mouse brain atlas (123.2*mm*^3^), this yielded a target number of 88.74 billion synapses. From this number we subtracted 23 billion synapses we predicted in local connectivity within a brain region, using previously described methods (Reimann et al., 2015). We then derived a matrix of synapse densities in all projections between pairs of brain region by scaling the “wild type connection density” matrices provided by Harris et al. (2018) in the following way:

Let *M_i_* and *M_c_* be the 43 ~ 43 matrices of connection densities in ipsi- and contra-lateral projections between brain regions, provided by Harris et al. (2018). Entries along the main diagonal of *M_i_*, corresponding to connectivity within a region are set to 0. Further let *V* be the vector of region volumes and *C_t_* the matrix of target region coverage in Fig. 7 D. Then we can calculate the scaling factor *σ*:

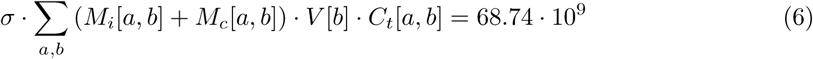

This factor was then applied to both *M_i_* and *M_c_* to convert them into matrices of the average density of synapses in the target region due to a projection, measured in *µm^−^*^3^ While this left no explicit room for synapses from extra-cortical sources, we estimate them to contribute comparatively little. For example, the density of thalamic synapses projected from VPM into SSp-bfd (Meyer et al., 2010), when averaged over the whole cortical depth, is only about 1.5% of the average total density (0.72*µm^−^*^3^, Schüz and Palm (1989)).

#### 6.1.1 Projection density matrices for individual projection types

We combined the “wild type” projection matrix from Harris et al. (2018) with their incomplete information on projections in individual projection classes, to get five individual projection matrices, one for each projection class. As their “wild type” experiments affected neurons in all layers and classes of the source region, we assumed that the sum of synapse densities over projection classes is equal to the density for the wild type. Further, based on qualitative observations, we assumed that the region-to-region connection matrices for each projection class are versions to the wild-type matrix, where individual *module-to-module submatrices* are scaled by individual values. The *modules* were six groups of contiguous brain regions (prefrontal, anterolateral, somatomotor, visual, medial, temporal) identified in Harris et al. (2018). This assumption means that connectivity trends between modules will be preserved for all projection classes but more fine grained trends for regions within a module will simply replicate the overall trends observed in the “wild type” matrix for all classes.

Based on these assumptions, we derived matrices of synapse densities for individual projection classes with the following algorithm. First, we digitized the available information for individual projection classes from the Harris paper and condensed it into five 6 ×6 matrices of average projection strengths between modules. We then normalized results such that the sum of the five matrices is 1 for each entry. Finally, we generated full-size 43 by 43 matrices for each projection type by scaling module-to-module specific submatrices of the wild type matrix by the corresponding entry in the condensed and normalized matrix (Fig. S1).

**Figure S1:**
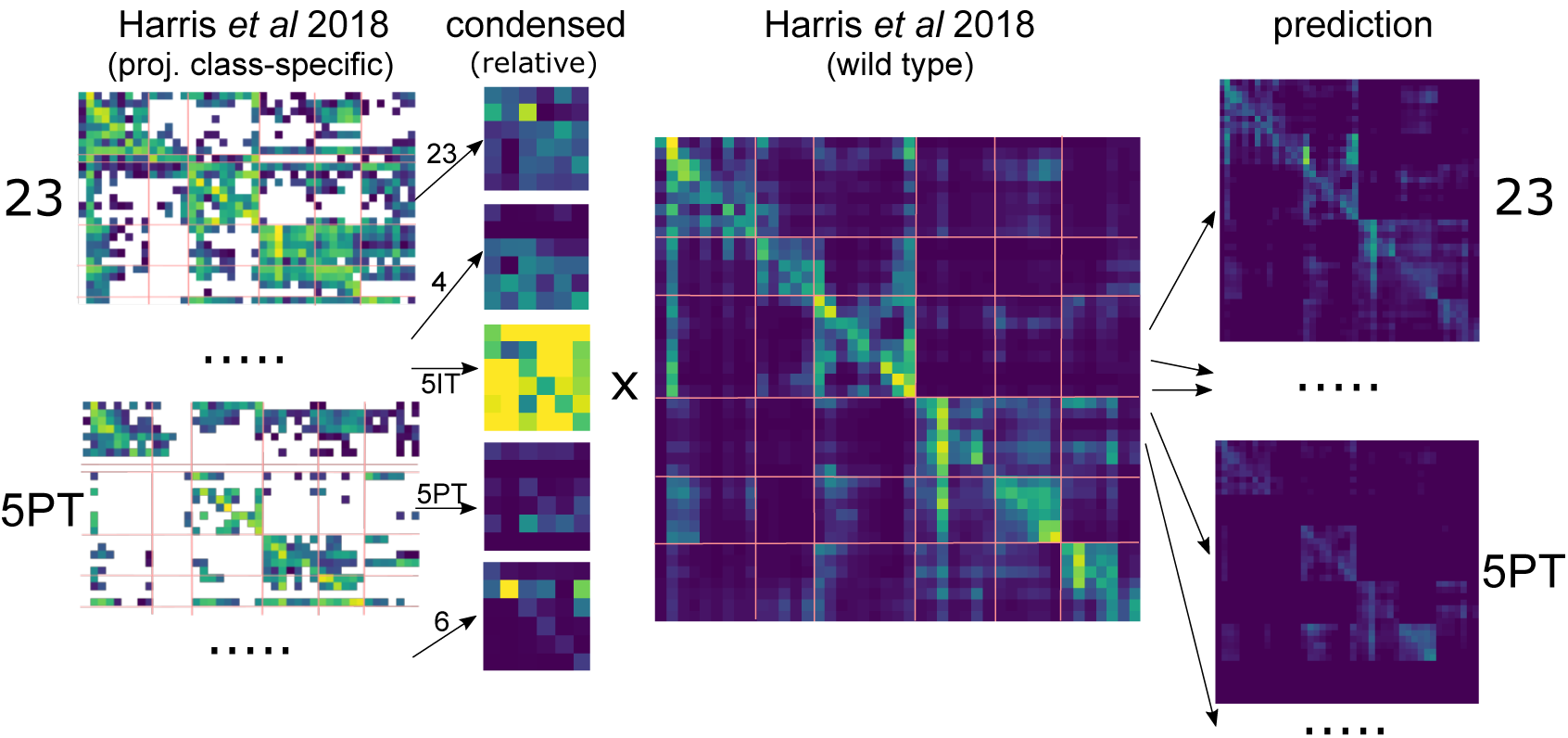
Prediction of 43 by 43 projection class-specific connection matrices

### 6.2 Predicting layer profiles

To assign one out of six layer profiles to each projection, we digitized the data on profile frequencies of Harris et al. (2018) and combined it according to the process illustrated in Fig. S2: First, for a source module we counted the number of intra-module or inter-module projections originating from it in each projection class. The example illustrates inter-module feed-forward projections from the prefrontal module (Fig. S2, top left). For the presence of a projection we defined a minimum projection strength, selected such that less than 5% of the total number of projection synapses are lost to the cutoff. The counts were then used as weights for a weighted average of the vectors of layer profile frequencies associated with each projection class. The result is a vector of *expected profile frequencies* for intra- or inter-module projections from the source module, if only the layer profile frequencies associated with projection classes are considered (Fig. S2, top right).

**Figure S2:**
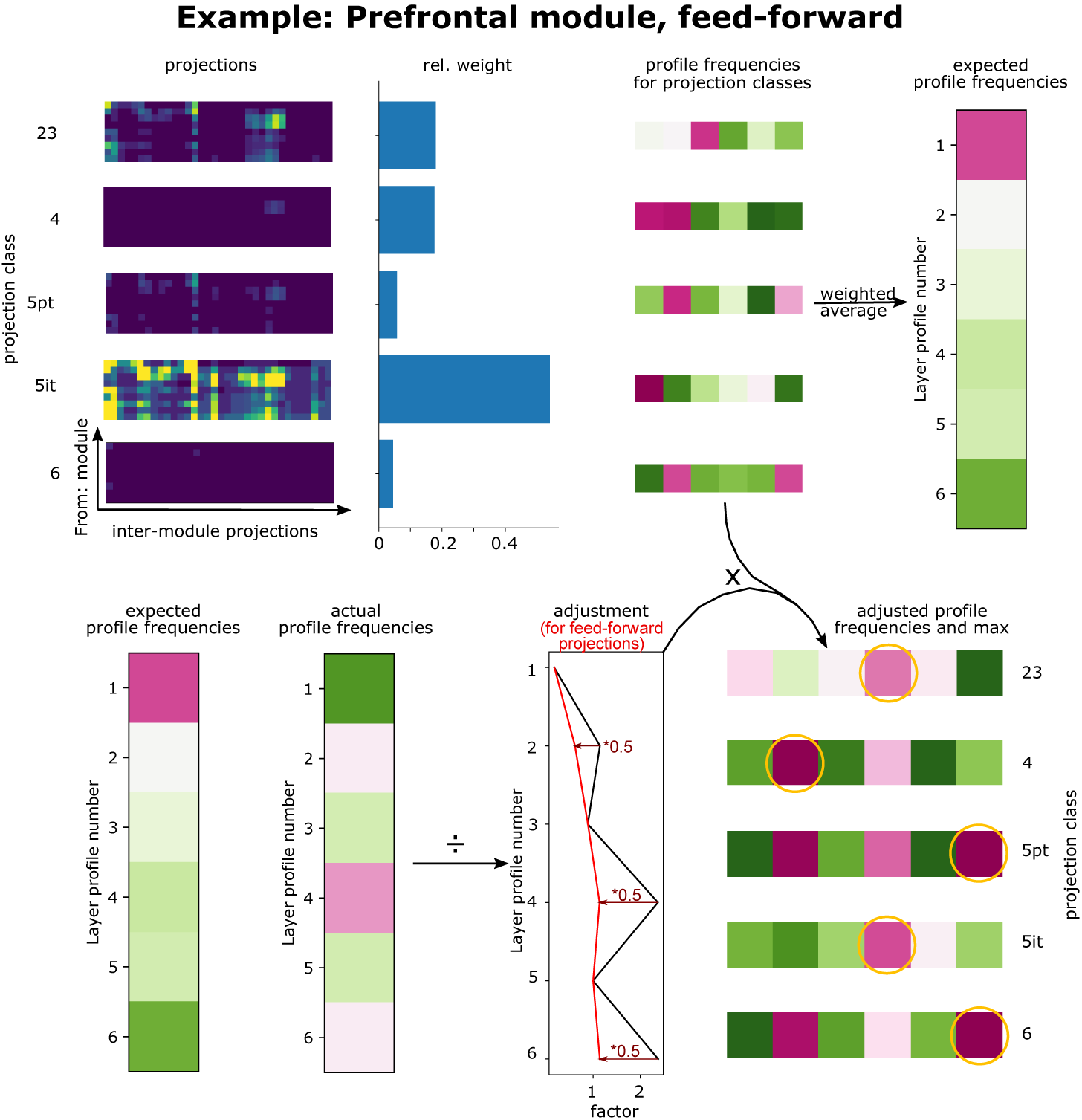
Predicting the layer profile frequencies for feed-forward inter-module connections from the prefrontal module and all projection classes

Next, we looked up the observed profile frequencies for the source module in the data of Harris et al. (2018) and compared them to the expected ones (Fig. S2, bottom left). Dividing the observed by the expected frequencies yielded adjustment factors for each layer profile that expressed which profiles were over- or under-expressed in intra- or inter-module projections from the source module under consideration (Fig. S2, bottom middle). We categorized projections as feed-forward or feed-back, based on the hierarchical positions of brain regions, reported in Fig. 8e of Harris et al. (2018), and, in accordance with their findings, reduced by 50% the adjustment factors for profiles 1, 3 and 5 when considering feed-back projections and of profiles 2, 4, 6 when considering feed-forward projections. Finally, we multiplied the vector of adjustment factors with the vectors of profile frequencies for individual projection classes to get *adjusted profile frequencies* (Fig. S2, bottom right).

The method yielded unique profile frequencies for each combination of source module, projection class and intra- or inter-module projection. To reduce the vectors of adjusted frequencies to a single profile, we simply picked the profile with the highest adjusted frequency (Fig. S2, bottom right).

### 6.3 Analyzing whole-brain axons

We acquired 183 neuron reconstructions from the Janelia Mouselight data portal (Gerfen et al., 2018) by querying for reconstructions where the soma location is within the neocortex. We first manually annotated the apical dendrite using Neurolucida (MBF Bioscience, Williston, VT USA) given that it was not available in the original data. Based on this we classified the neuron as a pyramidal cell or interneuron. Then, we performed a spatial analysis of the axon projection of each neuron by mapping the terminal points of the axon as well as the soma location into the Allen CCFv3 atlas coordinate system (Oh et al., 2014). This yielded a complete list of brain regions containing axon terminal branches, as well as the brain region and layer containing the soma. Together with the information previously extracted from the annotated apical dendrite (e.g. shape, layer, number of branches), this spatial information is used to perform classification of the m-type and projection type (p-type) of the neuron.

### 6.4 Constructing the p-type generating tree morphology

The Louvain algorithm takes a weighted adjacency matrix as input and then clusters the nodes into communities trying to maximize the weights within a community and minimize the weights across. An additional parameter is *γ*, which defines the granularity of the result: The smaller the value, the fewer communities it will result in, until a value of zero resulting in a single community.

We began by setting *gamma* to a value of 6.0, such that every brain region resulted in its own community. Correspondingly, we began constructing the tree topology by associating every brain region with its own leaf node. We then continuously lowered the value of *γ*, such that regions and communities began to merge into larger communities. We considered a pair of communities to be merged when through lowering *γ* a new community appeared that contained more than half of the regions of each of the original communities. In that case, we placed a new node in the graph representing the new community and connected it with the two nodes representing the original communities. We continued lowering *γ* until it reached zero, at which point everything merged into a single community and the root of the tree was placed.

We fit the weights of the edges to the predicted innervation probabilities using a recursive algorithm that optimized the local weights in small motifs consisting of two sibling nodes and their parent. It is based on the following observations (Figure S3):

**Figure S3:**
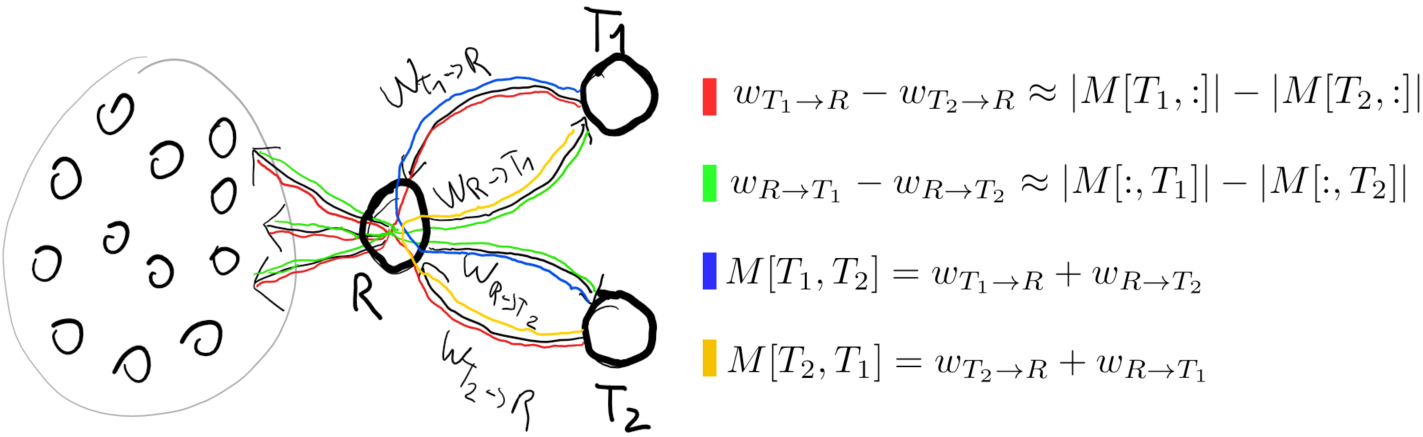
The weights in the p-type model tree can be estimated for triad motifs of two siblings and their parent from the matrix of path lengths between leaf nodes and four known shortest paths in the motif. Blue: shortest paths from leaf nodes to all other leaves; red: shortest paths from all other leaves to the leaf nodes; green and orange: shortest paths between the leaf nodes.

Let *T*_1_ and *T*_2_ be two sibling nodes and *R* their parent. In the model, any difference in the innervation probabilities for axons originating in *T*_1_ and *T*_2_ can only be due to differences in the lengths of the edges connecting each of them to their parent. This is because once the parent is reached, the shortest paths to any other region will be identical. Therefore:

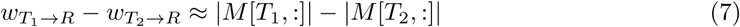

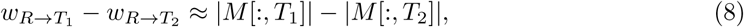

Where *M* denotes the matrix of the negative logarithm of predicted innervation probabilities, *M* [*x,*:] a single row of it (i.e. the probabilities of neurons in *x* to innervate each other region) and *M* [:*, x*] a single column of it (i.e. the probabilities of *x* to be innervated by neurons in each other region).

Further, the probability a neuron in *T*_1_ innervates *T*_2_ is given by the path from *T*_1_ via *R* to *T*_2_:

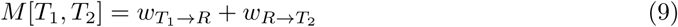

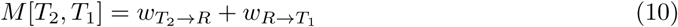

We found values for the four edge lengths in the motif by finding the least-squares solution of the system of linear equations. After this, we continued by performing the same step for node *R* and its sibling, until the root was reached.

### 6.5 Generating connectome instances according to the constraints

As mentioned previously, a *long-range projection recipe* is created which describes constraints on the desired connectivity. By doing so, the same recipe can be used to instantiate long-range projections with different circuit models, and to allow for different implementations to create these instantiations. This section describes the implementation used to generate the connectomes published under https://portal.bluebrain.epfl.ch/resources/models/mouse-projections.

The circuit representation and input data required for this implementation are:

- A placement of neuron morphologies in space
- A table describe their morphological types
- A spatial index, allowing the querying of morphology segments in a bounding region
- An atlas describing the different regions and layers that are addressed by the recipe
- A ‘flat-mapping’ from 3D to 2D space
- the recipe, which has:

– populations: Defining which regions, subregions and morphological types are part of the various source and target populations
– projections: organized by source population; specifies per target population, the expected synapse density, layer profile, and the barycentric source and target triangles
– p-types: organized by source population; specifies per target population the first order innervation probabilities for neurons of the source population, and for pairs of target populations the conditional increase in innervation probabilities

The basic circuit representation (first three items) was generated by a scaled-up version of a previous algorithm (Markram et al., 2015). The atlas was based on the Allen Common Coordinate Framework (Lein et al., 2006). For the flat-mapping we used the “Allen Dorsal Flatmap” of the “mcmodels” python package (see above). With this data, the implementation proceeds with the following steps:

#### Neuron allocation

For each source population, the neurons in those populations are allocated to participate in projections to a number of target populations according to specified fractions and statistical interactions. Where no interaction is specified, the overlap (neurons participating in both projections) is calculated from the fractions participating in one projection multiplied by the other; this default value is scaled up where interactions are specified. The challenge is then to assign neurons to each of the projections, such that the desired fractions and overlap sizes are reached.

A simplistic greedy algorithm was used to perform this allocation. Each source population group is assigned a sampled set of neurons, and pairwise the overlap is calculated, and adjusted based on the first order interactions. When the overlap is too small, it is enforced by randomly sampling neurons from each group, and replacing neurons in the other group such that the overlap is achieved. Attempts were made to use a SAT solver to perform exact allocations, but the size of the neuron counts and the constraint counts meant the model could not be solved in the available memory.

#### Synapse sampling

Sampling happens at the target region level. The target populations in the region are grouped, and the required densities per incoming projection are computed based on the long-range projections recipe. The densities are translated into counts based on the constrained volume created by intersecting the area occupied by the barycentric triangles, and reversed mapped using the ‘flat-map’ to the voxels of the atlas within the region. Finally, all morphological segments of a target population within this volume are found, and sampled with replacement with weights proportional to their length. Synapses are placed at random offsets within these segments. This structure allows for parallelization, as each combination of target and source populations can be run at the same time, subject to computation and memory limits. In practice, finding all segments within the volume demands significant memory which constrains the implementation. This, in turn, gives rise to per target population order: all samples for this population are loaded, and then all the sources referencing this population are calculated sequentially, with the calculations parallelized when possible - the initial sampling, picking the segments with in the barycentric coordinates, etc. Further parallelization can be achieved by running many of these process on different machines, in a batch style.

#### Mapping

Following the allocation and sampling, the two results are brought together in mapping: source neurons that are allocated to a projection are matched to the synapses created during the sampling of the same projection. Because both these datasets work with 3D coordinates, they are projected into the 2D representation so that the barycentric coordinates, described earlier, can be used to create the desired spatial organization.

To that end, since source neurons are less numerous, they are first projected into the flat space, and from there mapped into the barycentric coordinate system of the source region. The same coordinates in the barycentric system of the target region are then mapped back into the flat space and considered the *mapped locations* of the source neurons in the target region. Synapses in the target region are directly mapped to the flat space. Finally, in parallel, synapses are then stochastically assigned to a target neuron with a weighting based on the distance to their mapped location in the flat space and the specified width of the mapping. To speed this process, the source locations are put in a k-dimensional tree, and only the 100 closest source locations are queried per potential target synapses.

#### Output

The final step is to output the circuit in a format that can be used for simulation. For this, the SONATA file format was chosen - https://github.com/AllenInstitute/sonata.

Additionally, for structural analysis, we output for each target region a connectivity matrix of all incoming connections in the scipy.sparse.csc matrix format.

## 7 Supplementary material

**Table S1:** Order of brain modules (left) and regions (middle and right) used throughout the manuscript

|  |  |  |
| --- | --- | --- |
| Prefrontal | FRP | Frontal pole, cerebral cortex |
|  | MOs | Secondary motor area |
|  | ACAd | Anterior cingulate area, dorsal part |
|  | ACAv | Anterior cingulate area, ventral part |
|  | PL | Prelimbic area |
|  | ILA | Infralimbic area |
|  | ORBl | Orbital area, lateral part |
|  | ORBm | Orbital area, medial part |
|  | ORBvl | Orbital area, ventrolateral part |
| Anterolateral | AId | Agranular insular area, dorsal part |
|  | AIv | Agranular insular area, ventral part |
|  | AIp | Agranular insular area, posterior part |
|  | GU | Gustatory areas |
|  | VISC | Visceral area |
| Somatomotor | SSs | Supplemental somatosensory area |
|  | SSp-bfd | Primary somatosensory area, barrel field |
|  | SSp-tr | Primary somatosensory area, trunk |
|  | SSp-ll | Primary somatosensory area, lower limb |
|  | SSp-ul | Primary somatosensory area, upper limb |
|  | SSp-un | Primary somatosensory area, unassigned |
|  | SSp-n | Primary somatosensory area, nose |
|  | SSp-m | Primary somatosensory area, mouth |
|  | MOp | Primary motor area |
| Visual | VISal | Anterolateral visual area |
|  | VISl | Lateral visual area |
|  | VISp | Primary visual area |
|  | VISpl | Posterolateral visual area |
|  | VISli | Laterointermediate area |
|  | VISpor | Postrhinal area |
|  | VISrl | Rostrolateral visual area |
| Medial | VISa | Anterior area |
|  | VISam | Anteromedial visual area |
|  | VISpm | posteromedial visual area |
|  | RSPagl | Retrosplenial area, lateral agranular part |
|  | RSPd | Retrosplenial area, dorsal part |
|  | RSPv | Retrosplenial area, ventral part |
| Temporal | AUDd | Dorsal auditory area |
|  | AUDp | Primary auditory area |
|  | AUDpo | Posterior auditory area |
|  | AUDv | Ventral auditory area |
|  | TEa | Temporal association areas |
|  | PERI | Perirhinal area |
|  | ECT | Ectorhinal area |

